# Mucosal delivery of influenza antigens using a replication deficient adenovirus supports broadly reactive antibody responses and heterologous viral immunity in the respiratory tract of animals

**DOI:** 10.1101/2025.10.13.682194

**Authors:** Michael D. Schultz, Davide Botta, Aaron Silva-Sanchez, Isabella R. Cheatwood, Davies Kalange, Sarah M. Terapane, Aaron C. K. Lucander, Fen Zhou, Jobaida Akther, Guang Yang, Mary E. Cockerham, William O. Stover, John A. Hall, Dinesh Yadav, Douglas M. Fox, Francisco Dominguez, Sixto M. Leal, Jeremy B. Foote, Kevin S. Harrod, Troy D. Randall, Adrian E. Rice, Elizabeth S. Gabitzsch, Frances E. Lund

**Affiliations:** Department of Microbiology, The University of Alabama at Birmingham, Birmingham AL 35294 USA; Immunology Institute, Heersink School of Medicine, The University of Alabama at Birmingham, Birmingham, AL 35294, USA; Department of Medicine, Division of Clinical Immunology and Rheumatology, University of Alabama at Birmingham, Birmingham, AL 35294, USA; Undergraduate Immunology Program, University of Alabama at Birmingham, Birmingham, AL 35294, USA; Department of Biology, Franciscan University of Steubenville, Steubenville, OH 43952, USA; Undergraduate Cancer Biology Program, College of Arts and Sciences and Heersink School of Medicine, The University of Alabama at Birmingham, Birmingham, AL 35294, USA; Department of Biology, Mercyhurst University, Erie, PA 16546, USA; Department of Biomedical Engineering, The University of Alabama at Birmingham, Birmingham, AL 35294, USA; Department of Pathology, Division of Laboratory Medicine, University of Alabama at Birmingham, Birmingham, AL 35294, USA; Southeastern Biosafety Laboratory Alabama Birmingham (SEBLAB), The University of Alabama at Birmingham, Birmingham, AL 35294, USA; Department of Anesthesiology and Perioperative Medicine, University of Alabama at Birmingham, Birmingham, AL 35294, USA; ImmunityBio, Inc., Culver City, CA 90232, USA

**Keywords:** Respiratory tract immunity, mucosal B cells, broadly reactive antibodies, influenza A, Adenovirus

## Abstract

Systemically administered influenza vaccines provide strain-limited protection, while influenza infection of the respiratory epithelium supports development of lung resident memory B and T cells and more broadly reactive antibody responses. To test whether local antigen delivery is critical for establishing broad immunity, we directly compared respiratory tract and systemic delivery of influenza antigens to mice and hamsters using a replication-deficient adenovirus serotype 5 vector (Ad5[E1-,E2b-,E3-]). Both immunization routes elicited antibody responses in the lower respiratory tract and antigen-specific B and T cells in the draining lymph node. However, only intranasal immunization established lung-resident memory B and T cells, induced IgA responses in the upper respiratory tract directly at the site of viral entry and supported generation of IgA and IgG antibodies that bound antigenically drifted and distantly related influenza strains, including those of avian origin. Intranasal immunization accelerated viral clearance following heterologous virus challenge and was associated with limited pulmonary inflammation and fibrosis. Thus, intranasal immunization with the immunologically stealthy Ad5[E1-,E2b-,E3-] platform supports respiratory and systemic immunity to divergent influenza strains in the absence of overt lung immunopathology, suggesting that local antigen delivery may be key to development of more broadly protective “universal” flu vaccines.

## Introduction

Respiratory virus infections, including infection caused by the influenza A (flu) virus, result in substantial morbidity and mortality, despite the availability of seasonal vaccines [1–4]. This is due, at least in part, to the ongoing selection of new seasonal influenza virus variants and the periodic emergence of new reassortment viruses. Despite annual reformulation of the inactivated flu vaccine, in most years the vaccine is unable to prevent infection or transmission of these constantly emerging viral variants [5–9]. This is in contrast to flu infection, which is reported to induce a broader humoral immune response relative to immunization [10–15]. Therefore, there is a need to understand why natural infection appears more effective than vaccination in eliciting the broadly reactive immune responses that can provide protection against viral variants [16, 17].

In contrast to the standard inactivated flu vaccine, which is given via the intramuscular (i.m.) route, the flu virus naturally and specifically infects the respiratory epithelium [18]. The presence of virus-specific immune effectors at this mucosal surface is believed to be important for blocking viral entry, reducing viral replication, and limiting virus transmission [19–22]. In particular, it is thought that secretory IgA and tissue-resident memory populations in the lungs and airways are critical for providing frontline protection that vaccine-induced systemic immunity often fails to achieve [23–25]. A growing body of literature using samples from infected and/or vaccinated humans indicates that the mucosal immune response, elicited after infection, is regulated by different mechanisms and is functionally distinct from the immune responses that take place in secondary lymphoid tissues [18, 26, 27]. As just one example, intramuscular (i.m.) vaccination using SARS-CoV-2 Spike mRNA results in the development of circulating virus-specific cells and antibodies [28, 29]. However, i.m. immunization does not efficiently elicit Spike-specific T and B cells in the lung tissue or airways [28, 29] and Spike-specific IgA is rarely detected in the airways or saliva of vaccinated individuals who have no prior recent history of infection [29, 30]. In striking contrast, virus-specific memory B and T cells as well as antibodies, particularly virus-specific IgA Abs, can be measured in the lungs, airways and saliva of previously infected individuals [28–30]. Thus, respiratory infection in humans appears to elicit a specific mucosal response that can provide protection at the site of virus entry.

Given the technical difficulties associated with in-depth sampling of human immune cells in the mucosal upper and lower respiratory tracts (URT and LRT), rodent and other animal models have been used to better characterize the mucosal B cell and humoral immune response elicited by infection [26, 31, 32]. Similar to humans, flu infection but not systemic immunization induces the development of lung flu-specific memory B cells in mice [23, 33]. Parabiosis experiments reveal that the lung flu-specific memory B cells induced after infection are resident and non-recirculating [23, 33]. The data further show that the presence of antigen within the lung [34–36] and T cells making the inflammatory cytokine IFNγ [37] are required for the development and/or placement of the lung BRM cells. In turn, B cell intrinsic expression of the IFNγ-induced transcription factor T-bet is required for the development and maintenance of the lung flu-specific BRM subset [38] and is also needed for the rapid differentiation of these cells into mucosa-localized antibody-secreting cells (ASCs, [38]). Thus, the presence of antigen within the local mucosal environment and specific anti-viral inflammatory signals appear important for the establishment of immune cells and mediators that can protect at the site of infection.

Data in mice further show that the lung memory B cells elicited after infection exhibit a broad reactivity profile and can bind to multiple influenza hemagglutinin (HA) antigens [39]. These results, which are consistent with the data in humans showing that flu infection induces a broader humoral immune response compared to standard i.m. immunization [15], suggest that the respiratory tract B cells and ASCs are programmed within the specialized mucosal environment to make qualitatively different types of responses. However, the factors contributing to development of a broadly reactive antibody response within the mucosa are still largely unknown. Based on experiments in flu-infected mice, it is postulated that sustained local lung immune responses, driven by the continued presence of antigen and inflammatory mediators driven by the respiratory flu infection, might be important [39]. However, most experiments to date have compared responses following productive flu infection to responses induced after immunization using flu proteins or flu virus that are administered to non-mucosal sites, like the peritoneal cavity, the muscle or the skin. Since immunization with flu proteins or flu virus outside of the respiratory tract does not result in productive infection or viral replication, systemic vaccination and infection differ substantially in the amount of antigen present, the duration of antigen availability, and the cytokine microenvironment.

Given that the differences between mucosal active infection and systemic immunization with replication deficient antigens extend well beyond the location of the response, we asked whether the development of a broadly reactive local antibody response in the lung requires viral replication. To do this, we turned to replication deficient adenovirus (Ad) that can be modified to express genes of interest [40], can elicit strong humoral and cellular immune responses [41–43] and can be given via the mucosal or systemic route [44, 45]. In the present study, we employed the replication-deficient adenovirus serotype 5 (Ad5) platform with deletions in E1, E2b, and E3 genes (Ad5 [E1-, E2b-, E3-]) that dampen innate recognition and anti-vector responses [46–50]. Using the Ad5 [E1-, E2b-E3-] vector, which was engineered to express either flu HA [51] or flu nucleoprotein, we directly compared immune responses to the same dose of replication-deficient virus given via a systemic or intranasal (i.n.) mucosal route. Consistent with prior publications using other Ad vector platforms [44, 45], we observed that lung resident memory flu-specific B and T cell responses were induced only following i.n. immunization with the replication deficient Ad5 vectors. However, we also observed that the cross-reactive Ab response within the URT of immunized mice and hamsters was only induced following i.n. immunization. This i.n. elicited mucosal response was associated with better protection after challenge infection with homotypic or heterosubtypic viruses as measured by more rapid viral clearance in the URT and less lung damage. Thus, while production of broadly reactive Abs within the respiratory tract requires mucosal antigen delivery, it does not appear to be dependent on productive virus replication or an ongoing overt anti-viral inflammatory response. This suggests that the specialized mucosal respiratory tract environment rather than antigen load per se is s necessary for development and maintenance of the resident mucosal B cells that can respond to virus variants at the site of infection. These findings have direct implications not only for influenza vaccine design but also for the broader application of Ad5-based mucosal vaccines against respiratory pathogens.

## Results

### Intranasal immunization with Ad5-delivered flu antigens induces systemic and mucosal antigen-specific B cell, T cell and antibody responses

Previous studies from our group [49, 51] showed that systemic administration of the replication deficient Ad5[E1-,E2b-,E3-] (abbreviated as Ad5) vector expressing influenza A hemagglutinin (HA) antigens elicited strong systemic adaptive immune responses. Given the emerging data showing the importance of mucosal immunity in protection from respiratory viruses [18] and the data showing that infection rather than vaccination elicits a more broadly reactive antibody response [15], we evaluated whether the replication deficient Ad5[E1-,E2b-,E3-] (abbreviated as Ad5) vector platform could be used to induce a systemic and mucosal response following antigen delivery to the respiratory tract. We therefore immunized adult C57BL/6J (B6) mice via the i.n. route with 1×10⁹ viral particles (VP) of the empty Ad5 vector (Empty-Ad5) or with Ad5 expressing HA derived from the H1N1 A/California/04/2009 (CA09) virus strain (CA09HA-Ad5). Between day 7 (D7) and day 21 (D21) post-immunization, we isolated draining mediastinal lymph node (medLN) cells and performed flow cytometry using fluorochrome-labeled tetramers of CA09HA trimers to enumerate D7 CA09HA-specific (CA09HA^+^) plasmablasts (PB, CD19^+/lo^CD138^+^, Figure S1A), D21 CA09HA^+^ germinal center B cells (GCB, CD19^+^IgD^neg^CD38^lo^PNA^+^, Figure S1B) and D21 CA09HA^+^ memory B cells (MBC, CD19^+^IgD^neg^CD38^+^PNA^lo^, Figure S1B). By D7 post-immunization, we observed a significant increase in both the frequency (Figure S1A) and number (Figure 1A) of CA09HA^+^ PB in the CA09HA-Ad5 immunized mice relative to those receiving Empty-Ad5. Likewise, by D21 post-immunization, the frequency (Figure S1B) and number of CA09HA^+^ MBC (Figures 1B) and GCB (Figures 1C) were significantly elevated in animals receiving CA09HA-Ad5 compared to Empty-Ad5. Similar results were seen when we measured NP-specific B cell responses in the medLN following immunization with an Ad5 vector expressing influenza A nucleoprotein (NP-Ad5, Figure S2A-E). Thus, mucosal i.n. administration of antigens delivered using an Ad5-based vector system elicits systemic PB and B cell responses in secondary lymphoid tissue.

**Figure 1.**
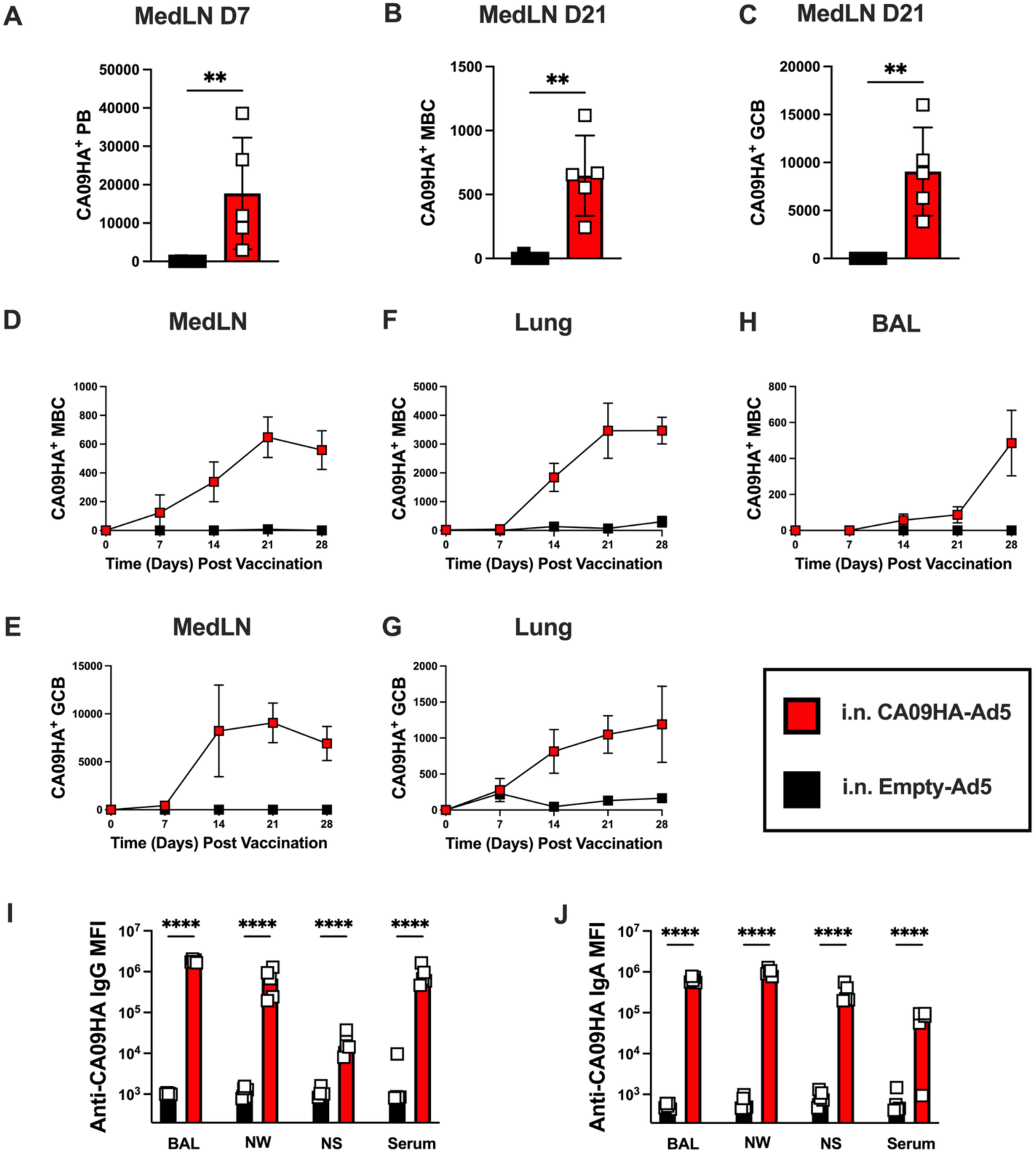
Intranasal immunization with CA09HA-Ad5 induces systemic and mucosal CA09HA-specific antibody and B cell responses. C57BL/6J (B6) mice (n=5 mice/group/timepoint/tissue sample) were immunized i.n. with Empty-Ad5 (black) or CA09HA-Ad5 (red). Samples (lung, mediastinal lymph node (MedLN), serum, bronchial alveolar lavage (BAL), nasal wash (NW), and nasal swipe (NS)) were collected at the indicated times and analyzed using flow cytometry (**A-G**) or cytometric bead arrays (CBA, **I-J**). (**A-C**) Quantitation of CA09HA-specific (CA09HA^+^) plasmablasts (PB, **A**), memory B cells (MBC, **B**) and germinal center B cells (GCB, **C**) in the MedLN on D7 (**A**) or D21 (**B-C**) following immunization. Data shown as mean ± SD of the medLN populations. (**D–H**) CA09HA^+^ MBC (**D, F, H**) and GCB (**E, G**) in the MedLN (**D-E**), lung (**F-G**) or BAL (**H**) between D0 to D28 post-immunization. Data shown as mean ± SEM of cells/tissue/timepoint. (**I–J**) CA09HA-specific Ab responses in serum, BAL, NW, and NS samples collected at D30 post-immunization. CA09HA-specific IgG (**I**) and IgA (**J**) levels were measured by CBA and reported as the mean ± SD of the log transformed mean fluorescence intensity (MFI) values of IgG or IgA Ab binding to CA09HA-coupled microbeads (see Methods). Representative flow cytometry gating for CA09HA^+^ PB, MBC and GCB in each tissue shown in Figure S1. Examination of NP^+^ B cell subsets following i.n. immunization with NP-Ad5 shown in Figure S2. Data analyzed using unpaired Mann–Whitney test (**A-C**) or two-way ANOVA with Tukey’s multiple comparisons test (**I-J**). * p < 0.05, ** p < 0.01, *** p < 0.001, **** p < 0.0001.

To address whether immunization with CA09-HA via the i.n. route also induced mucosal B cell responses, we used flow cytometry to enumerate CA09HA^+^ GCB and MBC in medLN (Figure S1B), lung (Figure S1C) and bronchial alveolar lavage (BAL, Figure S1D) samples collected between D7-D28 post-immunization from CA09HA-Ad5 and Empty-Ad5 immunized mice. As expected, CA09HA^+^ MBC (Figure 1D) and GCB (Figure 1E) were detected in the medLN. These cells could be observed as early as D14, peaked by D21 and remained elevated through at least D28 post-immunization (Figure 1D-E). Similarly, CA09HA^+^ MBC and GCB (Figure 1F-G) were detected in the lung by D14 with the number of CA09HA^+^ MBC continuing to increase through at least D28 (Figure 1F). CA09HA^+^ MBC were also detected in the BAL by D28 (Figure 1H).

Next, we asked whether flu antigen-specific antibodies (Abs) could be detected at systemic and mucosal sites following i.n. delivery of CA09HA-Ad5 or NP-Ad5. Therefore, we immunized mice with Empty-Ad5 and either CA09HA-Ad5 or NP-Ad5 via the i.n. route. On D28 we collected serum that contained the systemic Abs, BAL that contained Abs present in the lung airways and lower respiratory tract (LRT), nasal wash (NW) that contained Abs present in the upper respiratory tract ((URT), oral pharyngeal and nasal cavities) and nasal swipes (NS) that contained Abs present at the interface between the nasal cavity and the outside environment. Samples were incubated with cytometric beads arrayed (CBA) with recombinant CA09HA or flu NP proteins. Binding to CA09HA antigen by the polyclonal Abs present in each sample was detected using fluorochrome-labeled secondary Abs specific for murine IgG or IgA. Consistent with the detectable systemic B cell response in the medLN, CA09HA-specific IgG was detected in serum as well as NS, NW and BAL samples (Figure 1I). CA09HA-specific IgA responses in the mucosal BAL, NW and NS samples were also significantly elevated in the CA09HA-Ad5 immunized animals compared to Empty-Ad5 immunized mice (Figure 1J). Similar results were observed in mice immunized with NP-Ad5 (Figure S2F-G).

Since i.n. immunization with the CA09HA-Ad5 elicited both mucosal and systemic B cell and Ab responses, we predicted that systemic and mucosal T cell responses would also be elicited by i.n. delivery of the Ad5 vector. To test this, we immunized B6 mice i.n. with NP-Ad5 or Empty-Ad5 and used T cell NP tetramers (H-2Db NP_366-374_) to monitor the frequencies (Figure 2A-C) and numbers (Figure 2D-F) of NP^+^ CD8⁺ T cells in the medLN (Figure 2A, D), lung (Figure 2B, E) and BAL (Figure 2C, F) between D7-28 post-immunization. Similar to the NP-Ad5-elicited B cell response, the NP^+^ CD8 T cell response was observed by D14 in both the medLN and mucosal sites and was maintained for at least 28 days post-immunization. Moreover, the frequencies (D21, Figure 2A-C) and absolute numbers (D14-D28, Figure 2D-F) of NP^+^ CD8⁺ T cells were significantly greater in all three tissues of the NP-Ad5 immunized animals compared to the Ad5-empty immunized mice. Taken together, these data show that i.n. administration of either CA09HA-Ad5 or NP-Ad5 elicits systemic and respiratory tract cellular and humoral immunity.

**Figure 2.**
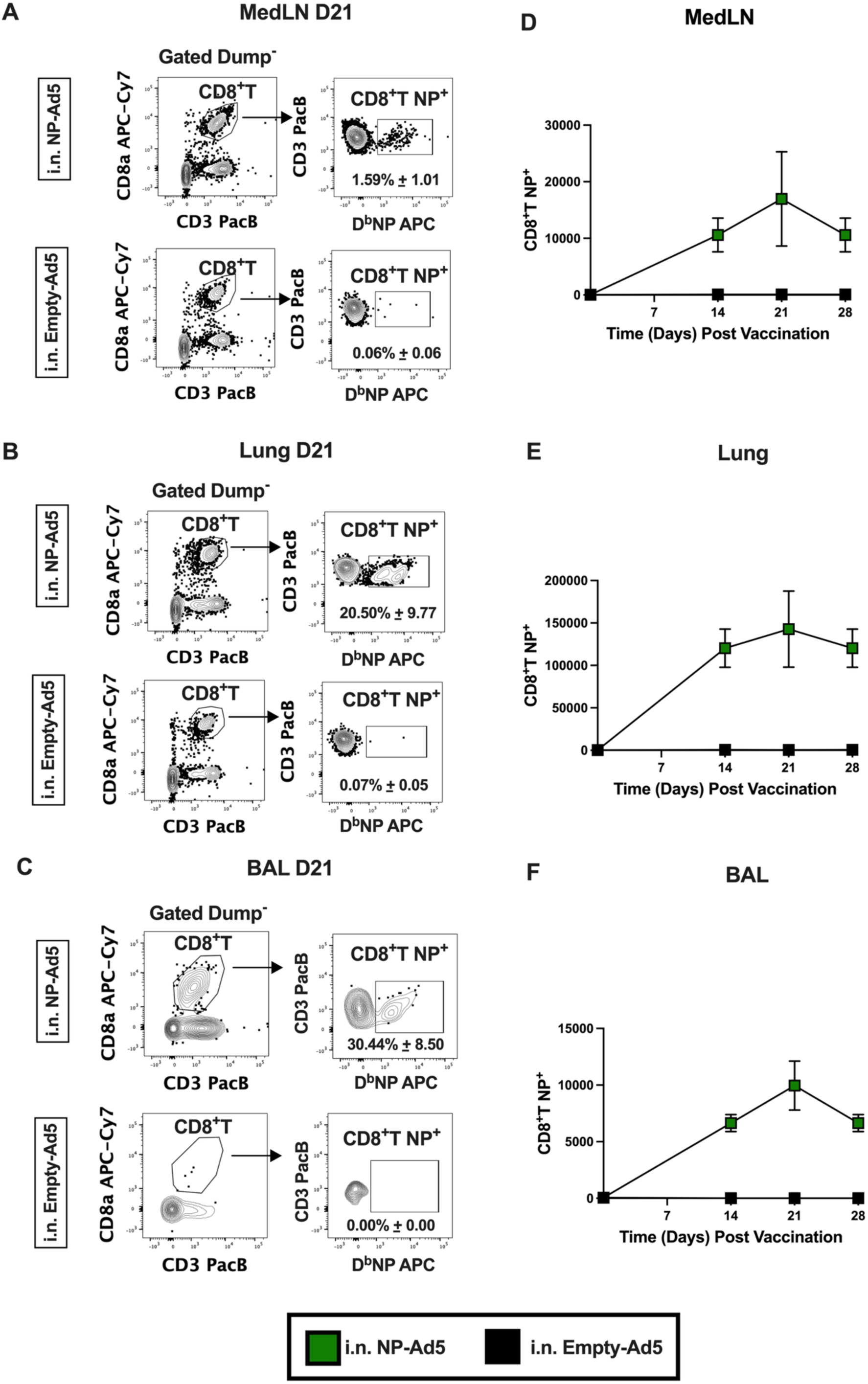
Intranasal immunization with NP-Ad5 elicits systemic and mucosal NP-specific T cell responses. B6 mice (n=5 mice/group/timepoint/tissue sample) were immunized i.n. with Empty-Ad5 (black) or NP-Ad5 (green). MedLN, lung and BAL samples were collected at the indicated timepoints post-immunization and analyzed using flow cytometry. (**A-C**) Representative flow cytometry gating strategies for identification of NP-specific (H2Kb NP_366-374_) CD8⁺ T cells in live, Dump^neg^ (CD11b^-^CD64^-^) gated cells from MedLN (**A**), lung (**B**), and BAL (**C**) on D21 post-immunization. Population frequencies reported as mean ± SD. (**D-F**) Quantitation of NP-specific CD8⁺ T cells in the MedLN (**D**), lung (**E**), and BAL (**F**) between D0 to D28 post-immunization. Data reported as mean ± SEM of cells/tissue/timepoint.

### Intranasal immunization with CA09HA-Ad5 promotes accumulation of antigen-specific lung tissue and airway-residing B cells

We previously demonstrated that natural infection with influenza induces a population of T-bet-expressing, non-recirculating, lung resident MBC (B resident memory cells or BRM) that are localized within the lung parenchyma and airways [33, 38, 52]. We further showed B cell intrinsic expression of T-bet is required for the “effector” lung BRM to rapidly differentiate *in situ* into secondary PB that provide local protective immunity following challenge infection [38]. However, these T-bet^+^ effector MBC were not induced when influenza virus was administered intraperitoneally (i.p.) [33], suggesting that establishment of this protective MBC compartment requires local antigen delivery. To test whether i.n. delivery of CA09HA-Ad5 was sufficient to generate the effector T-bet^+^ BRM compartment, we administered CA09HA-Ad5 i.n. to either B6 mice or *Tbx21*-ZsGreen mice that express the fluorescent reporter ZsGreen under the control of the T-bet gene (*Tbx21*) promoter. We then used flow cytometry to detect CA09HA^+^ MBC expressing the T-bet reporter in the lung and BAL of D28 immunized animals. We observed significantly elevated frequencies (Figure 3A-B) and absolute numbers (Figure 3C) of T-bet^+^ HA tetramer-binding memory B cells in the lungs 28 days after immunization. Similarly, *Tbx21* reporter expressing MBC were easily detected and significantly elevated in the BAL of i.n. immunized mice (Figure 3D-F). Moreover, 75-85% of the CA09HA^+^ MBC found in lung (Figure 3B) and BAL (Figure 3E) expressed the *Tbx21* reporter. Thus, the majority of mucosal MBC elicited by i.n. administration of CA09HA-Ad5 resided in the respiratory tract and exhibited an effector BRM phenotype.

**Figure 3.**
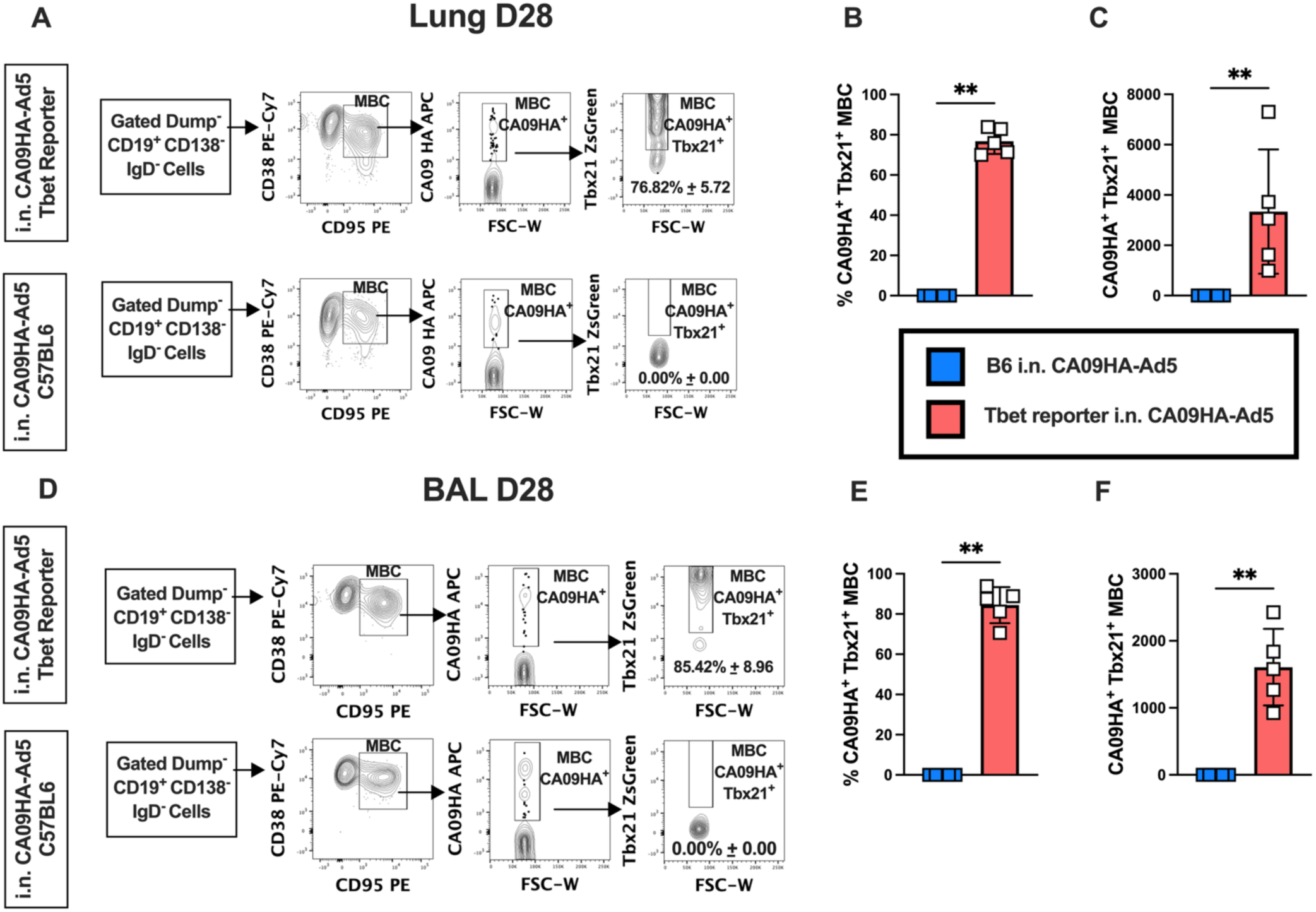
Intranasal immunization with CA09HA-Ad5 promotes establishment of flu-specific lung residing B cells. B6 (blue symbol) and T-bet reporter (*Tbx21*-ZsGreen, red symbol) mice (n=5 mice/group/timepoint/tissue sample) were immunized i.n. with CA09HA-Ad5. Lung and BAL samples were evaluated on D28 post i.n. immunization and analyzed using flow cytometry. Representative flow cytometry gating strategies to identify *Tbx21*⁺ (ZsGreen^+^) CA09HA^+^ MBC in lung (**A**) and BAL (**D**) are shown with the frequency (**A-B, D-E**) and number (**C, F**) of CA09HA^+^ *Tbx21*^+^ lung resident MBC reported as mean ± SD of cells/tissue. Data (**B-C, E-F**) analyzed using an unpaired Mann-Whitney test. * p < 0.05, ** p < 0.01, *** p < 0.001, **** p < 0.0001.

### Mucosal, but not systemic, administration of the Ad5-vectored immunogen induces antigen-specific lung memory B and T cells and promotes mucosal antibody responses

Our data indicated that i.n. immunization with CA09HA-Ad5 induced systemic, as well as local mucosal B cell immune responses. Given these results, we predicted that i.n. immunization with NP-Ad5 should also generate resident memory CD8^+^ T cells (CD8 TRM), which are known to support local rapid killing of virus infected cells. To test this, we immunized 6–8-week-old B6 mice with 1×10⁹ VP of Empty-Ad5 given i.n. or NP-Ad5 given either i.n. or i.p. and then used flow cytometry (Figure S3A-B) to enumerate the NP-specific CD8^+^ T cell subsets in medLN, which drains the peritoneal cavity and the lung. Few NP^+^ TRM (CD103^+^CD69^+^, Figure 4A) or NP^+^ memory (CXCR6^+^CXCR3^+/-^, Figure 4B) CD8 T cells were identified in the medLN at this timepoint, regardless of whether the animals were immunized via the i.n. route or i.p. route. However, NP^+^ effector (CXCR6^neg^CXCR3^+^) CD8 T cells were elevated in the medLN of the NP-Ad5 immunized mice (Figure 4C), with the largest expansion seen following i.n. immunization. Consistent with prior reports showing that lung CD8 TRM are generated following i.n. infection or i.n. immunization [53, 54], we identified a large population of CD8 TRM in the lung only after i.n. immunization (Figure 4D). Similarly, the NP^+^ CXCR6^+^CXCR3^+/-^ memory CD8 T cells were only expanded in the lungs of i.n. immunized mice (Figure 4E) while NP^+^ effector CD8 T cells were observed in the lungs of animals immunized by either route (Figure 4F). Therefore, the data showed that development of the CD8 TRM lung population is highly dependent on the NP-Ad5 vector being delivered intranasally.

**Figure 4.**
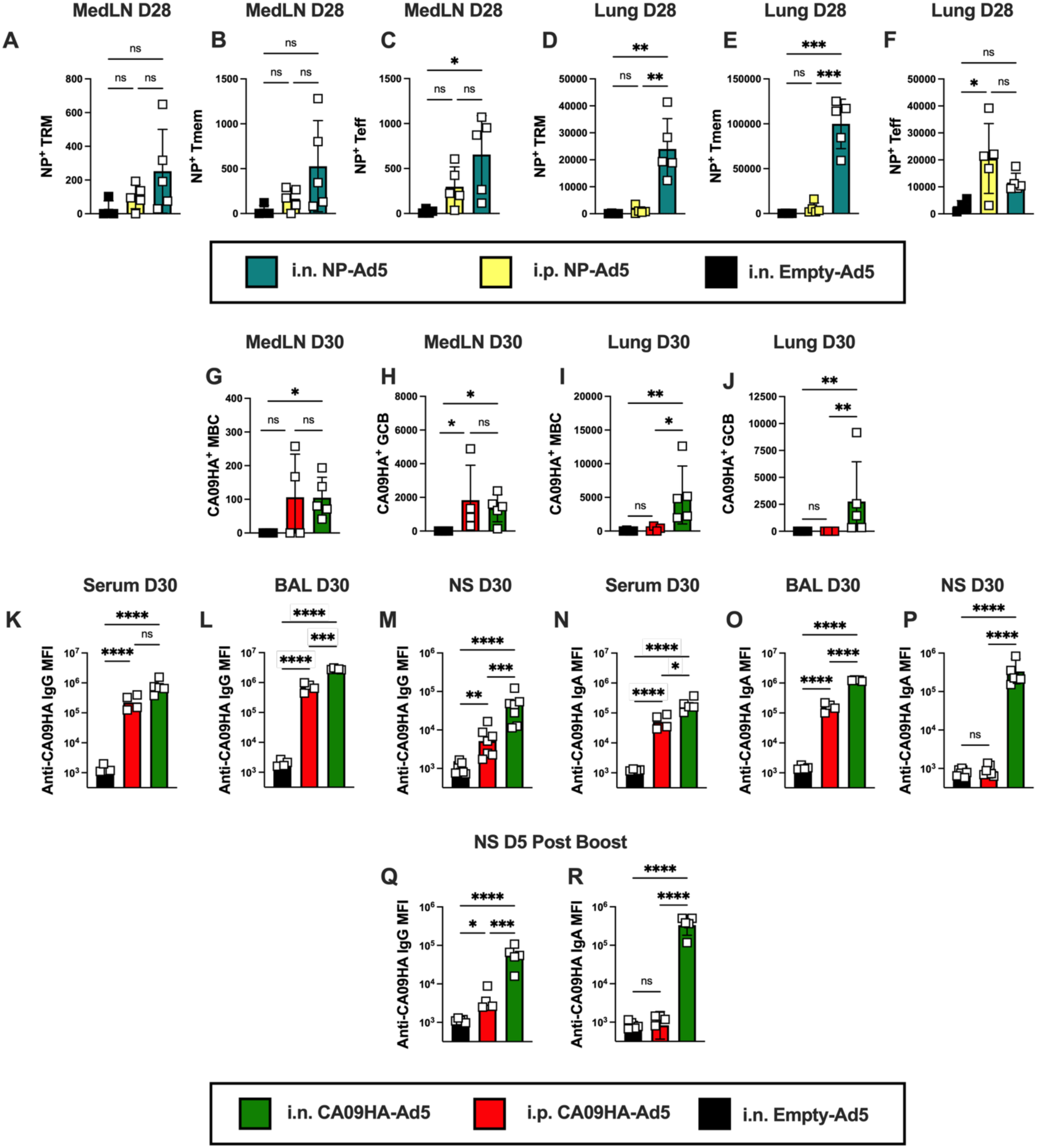
Mucosal, but not systemic, immunization with CA09HA-Ad5 establishes humoral immunity in the upper respiratory tract. B6 mice (n=4-7 mice/group/tissue sample) were immunized i.n. with Empty-Ad5 (black, all panels); were immunized with NP-Ad5 (panels **A-F**) via the i.n. (teal) or i.p. (yellow) route; or were immunized with CA09HA-Ad5 (**G-R**) via the i.n. (green) or i.p. (red) route. An additional cohort of animals (**K-L**, n=5 mice/group) was boosted on D30 with CA09HA-Ad5 by the same route (i.n./i.n. or i.p./i.p.). MedLN and lung tissues were collected on D28 (**A-F**) or D30 (**G-J**) for flow cytometry analysis. Serum, BAL, NW and NS samples were collected on D30 and analyzed by CBA (**K-P**). NS samples were collected on D5 post-boost (**Q-R**) and analyzed by CBA. (**A-F**) Number of NP^+^ CD69^+^CD103^+^ CD8 TRM (**A, D**), NP^+^ CXCR6^+^CXCR3^+/-^ CD8 Tmem (**B, E**) and NP^+^ CXCR3^+^CXCR6^-^ CD8 Teff (**C, F**) in the medLN (**A-C**) and lung (**D-F**) on D28 post-immunization. Data reported as mean ± SD for 5 mice/group/tissue sample. (**G-J**) Number of CA09HA^+^ MBC (**G, I**) and GCB (**H,J**) in the MedLN (**G-H**) and lung (**I-J**) on D30 post-immunization. Data reported as mean ± SD for 4-5 mice/group/tissue sample. (**K-P**) Quantitation of CA09HA-specific IgG (**K-M**) and IgA (**N-P**) in serum (**K, N**, n=4-5 mice/group), BAL (**L, O**, n=4-5 mice/group), and NS samples (**M, P**, n=7 mice/group) on D30 post-immunization. Data reported as mean ± SD of log transformed MFI values of Ab binding to CA09HA-coupled microbeads. (**Q-R**) Quantitation of CA09HA-specific IgG (**Q**) and IgA (**R**) in NS samples (n=5 mice/group) on D5 post-boost. Data reported as mean ± SD of log transformed MFI values of Ab binding to CA09HA-coupled microbeads. Representative flow cytometry gating to identify NP^+^ CD8 T cell subsets and CA09HA+ B cell subsets in lung and medLN shown in Figure S3. Data analyzed using two-way ANOVA with Tukey’s multiple comparisons test (**A-F, K-R**), or the Kruskal-Wallis test (**G-J**) * p < 0.05, ** p < 0.01, *** p < 0.001, **** p < 0.0001.

To address whether the establishment of the lung B cell and mucosal Ab response was also dependent on i.n. delivery of CA09HA-Ad5, we immunized 6–8-week-old B6 mice with 1×10⁹ VP of Empty-Ad5 given i.n. or CA09HA-Ad5 given either i.n. or i.p. On D30 post-immunization, lung, MedLN, serum, BAL and NS samples were collected for analysis. CA09HA^+^ GCB and MBC in lung and MedLN (Figure S3C-D) were detected using flow cytometry and CA09HA-specific IgG and IgA Abs in serum, BAL and NS were measured using CBA. Not surprisingly, the medLN harbored CA09HA^+^ MBC and GCB following immunization (Figure 4G-H) and there was no significant difference in the number of medLN CA09HA^+^ MBC and GCB between the i.n. and i.p. immunized groups. By contrast, CA09HA^+^ MBC and GCB were only detected in the lungs of animals immunized via the i.n. route (Figure 4I-J). Thus, i.p. delivery of CA09HA-Ad5 was not sufficient to support establishment of lung-residing MBC and GCB populations.

Both routes of administration induced a robust systemic serum CA09HA-specific IgG Ab response (Figure 4K) and there was no significant difference between the two immunized groups of mice. Likewise, CA09HA-specific IgG Abs were detected in BAL (Figure 4L) and NS (Figure 4M) samples from both immunized groups. However, the levels of CA09HA-specific IgG in BAL and NS samples were modestly but significantly higher in the i.n. immunized mice compared to the i.p. immunized animals (Figure 4L-M). Both immunization routes also generated CA09HA-specific IgA responses in serum (Figure 4N) and BAL (Figure 4O), but again the response was significantly elevated in animals immunized via the i.n. route. Strikingly, CA09HA-specific IgA in nasal secretions from the NS sample was only detected in the i.n. immunized mice with CA09HA-specific IgA in i.p. immunized mice at background (Figure 4P). Thus, only mucosal delivery of CA09HA-Ad5 elicited production of secretory IgA at the physiologic site of influenza virus entry that is located at the interface between the nostrils and the environment.

Given that a single systemic immunization with CA09HA-Ad5 was unable to induce an URT IgA response, we asked whether boosting systemically could elicit IgA in nasal secretions. We therefore primed mice with CA98HA-Ad5 given either i.n. or i.p. and administered a homologous boost (i.n.-i.n. or i.p-i.p.) on D30 post primary immunization. We then assessed the URT IgG and IgA CA09HA-specific Ab response in NS D5 post-boost in both groups of mice (Figure 4Q-R). Again, animals primed and boosted via the i.n. route had significantly increased levels of IgG present in the NS sample relative to the i.p.-i.p. group (Figure 4Q). Moreover, CA09HA-specific IgA in NS was only detected the i.n.-i.n. group (Figure 4R). Taken together, these data argue that local antigen exposure is needed to establish the lung resident memory B and T cells and the URT IgA Ab response.

### Immunization with CA09HA-Ad5 via the i.n. route uniquely supports a broadly reactive Ab response to influenza HA antigens

A prior study showed that infection of mice with influenza virus induces a broadly reactive HA response [55]. To address whether a broadly reactive Ab response was elicited following immunization with the CA09HA-Ad5 vector via the i.n. route, we compared the humoral immune response in 6-8-week-old B6 animals immunized i.n. to animals immunized via the i.m. route that is commonly used in humans [56]. For a negative control, mice were immunized i.n. with Empty-Ad5 and for a positive control, mice were infected i.n. with a non-lethal dose (500 PFU) of the non-mouse adapted H1N1 CA09 virus strain. NS samples were collected weekly from immunized and infected animals. On D30 post-immunization or infection, animals were euthanized, and serum, NW, and BAL were collected (Figure 5A). To assess the breadth of the humoral immune response, samples were analyzed using the multiplexed CBA platform arrayed with a panel of HA antigens that included H1_HA and H3_HA antigens derived from different seasonal/pandemic viral strains, as well as H2 _HA and H5_HA antigens derived from avian viral strains.

**Figure 5.**
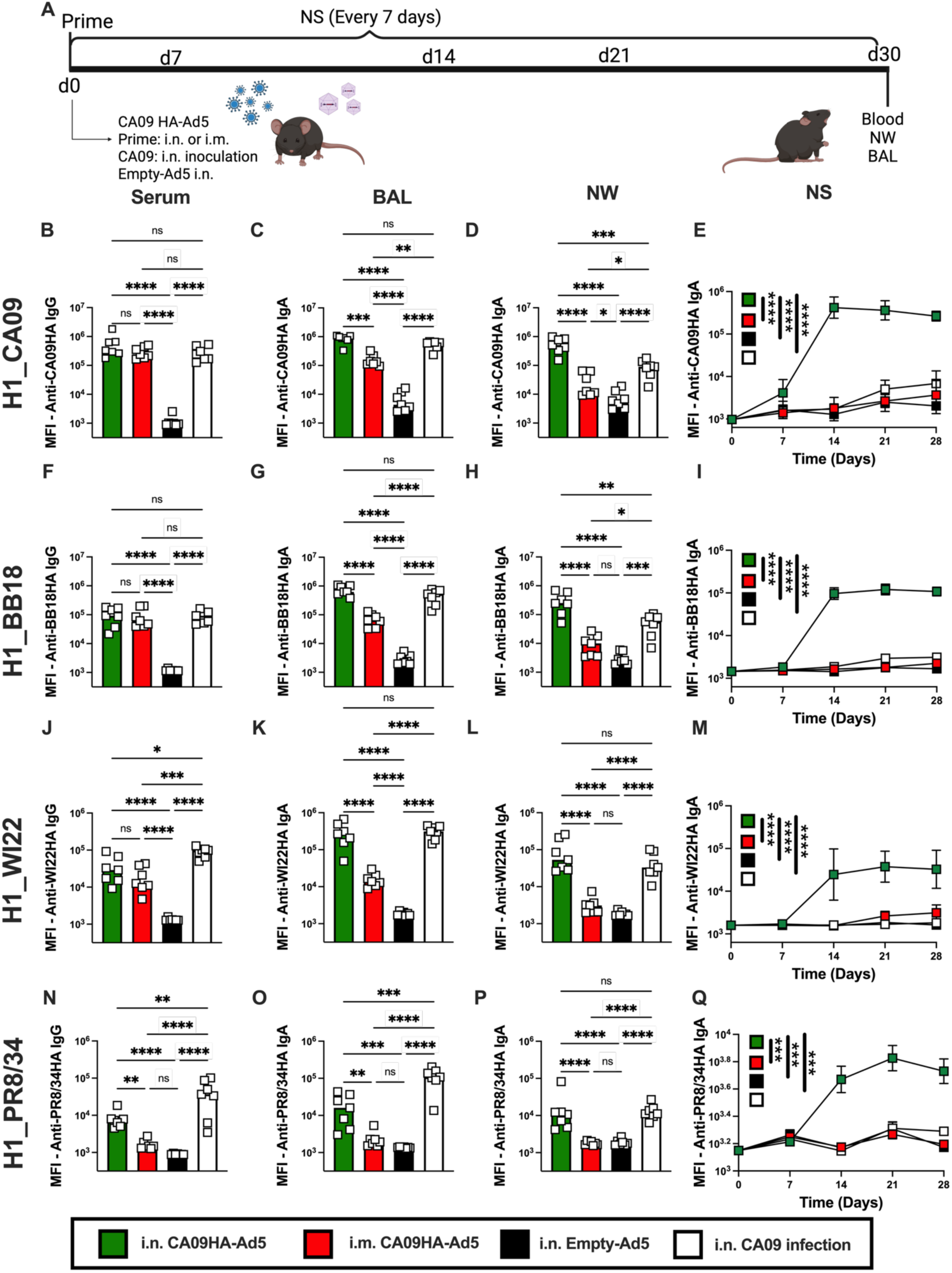
Intranasal immunization with CA09HA-Ad5 uniquely supports broadly reactive Ab responses in the upper respiratory tract. B6 mice (n=7/group) were infected i.n. with 500 plaque-forming units (PFU) H1_CA09 virus (white) or were immunized i.n. with Empty-Ad5 (black) or CA09HA-Ad5 via the i.n. (green) or i.m. (red) route. Samples were collected weekly (NS) or on D30 post-immunization/infection (serum, BAL, NW). Ab levels in samples were quantitated by CBA using microbeads coupled to recombinant H1_CA09HA, H1_BB18HA, H1_WI22HA, or H1_PR8/34HA antigens. (**A**) Study design schematic. (**B-E**) Quantitation of H1_CA09HA-binding IgG in serum (**B**) and H1_CA09HA-binding IgA in BAL (**C**), NW (**D**), and NS samples (**E**). (**F-I**) Quantitation of H1_BB18HA-binding IgG in serum (**F**), and H1_BB18HA-binding IgA in BAL (**G**), NW (**H**), and NS samples (**I**). (**J-M**) Quantitation of H1_WI22HA-binding IgG in serum (**J**), and H1_WI22HA-binding IgA in BAL (**K**), NW (**L**), and NS samples (**M**). (**N-Q**) Quantitation of H1_PR8/34HA-binding IgG in serum (**N**), and H1_PR8/34HA-binding IgA in BAL (**O**), NW (**P**), and NS samples (**Q**). Data reported as mean ± SD (**B-D, F-H, J-L, N-P**) or mean ± SEM (**E, I, M, Q**) of log transformed MFI values of Ab binding to the different HA-coupled microbeads. Quantitation of BAL, NW, NS IgG Ab binding to H1_CA09HA, H1_BB18HA, H1_WI22HA and H1_PR8/34HA shown in Figure S4. Quantitation of serum, BAL, NW, NS IgG and IgA Ab binding to H3_DW21, H2_NL99 and H5_VN04 shown in Figure S5. Statistical analysis performed using two-way ANOVA with Tukey’s multiple comparisons test (**B-D, F-H, J-L, N-P**) or two-way ANOVA on area under the curve (AUC) measurements (**E, I, M, Q**). * p < 0.05, ** p < 0.01, *** p < 0.001, **** p < 0.0001.

As expected, mice immunized with CA09HA-Ad5 or infected with CA09 virus developed comparable levels of serum H1_HACA09-specific IgG (Figure 5B) and this response was several orders of magnitude higher than seen in serum from animals immunized with Empty-Ad5. Consistent with our earlier experiment, i.n. immunized animals exhibited superior CA09HA-specific IgG (Figure S4A) and IgA (Figure 5C) responses in BAL when compared to i.m. immunized mice. In fact, the IgA BAL response in the i.n. immunized animals was equivalent to that seen in the infected mice (Figure 5C), indicating that i.n. delivery of CA09HA-Ad5 was as effective as influenza infection in eliciting a lung mucosal IgA response. The CA09HA-specific IgA (Figure 5D) and IgG (Figure S4B) responses in NW of i.m. immunized mice were not substantively higher than those detected in the Empty-Ad5 immunized mice. By contrast, the i.n. immunized mice made robust IgG (Figure S4B) and IgA (Figure 5D) responses in the URT (NW) that were comparable or even greater than those seen in the NW from CA09-infected mice. In addition, the CA09HA Ab response in the NS, which captures Abs at the interface of the nasal cavity with the environment, was largely limited to i.n. immunized mice (Figure S4C, Figure 4E). While low levels of CA09HA-specific IgG were detected in NS samples (Figure S4C), the major isotype present in that site was IgA (Figure 4E). Consistent with published data showing that replication of non-mouse adapted flu virus in the URT of wild-type mice is limited, particularly in the nasal cavity and turbinates [47], CA09HA-specific IgA was not observed in NS samples from infected mice and was exclusively detected in the i.n. immunized animals (Figure 5E).

Next, we examined whether IgG and IgA elicited following CA09HA-Ad5 immunization or CA09 infection exhibited binding to HA antigens from seasonally drifted strains of the 2009 H1_CA09 virus, including H1_BB18 (Figure 5F-I, Figure S4D-F), which circulated in 2018, and the H1_WI22 (Figure 5J-M, Figure S4G-I), which circulated in 2022. Again, IgG that exhibited reactivity for these seasonal drifted H1_HA antigens were present in serum (Figure 5F, 5J). Likewise, BAL IgA (Figure 5G, 5K) and BAL IgG (Figure S4D,G) capable of recognizing the related H1_HA antigens were readily detected in the immunized and infected mice, although the response was significantly lower in the i.m. immunized mice relative to the other groups. However, the amount of IgA (Figure 5H, 5L) and IgG (Figure S4E, S4H) in NW samples that could bind the drifted HA antigens were not different between the negative control, Empty-Ad immunized animals and the mice immunized i.m. with CA09HA-Ad5. By contrast, the i.n. immunized and flu-infected mice generated equivalent and high levels of NW IgA (Figure 5H, 5L) and IgG (Figure S4E, S4H) that bound the drifted H1_HA antigens. Moreover, we observed minimal IgA (Figure 5I, 5M) and IgG (Figure S4F, S4I) reactivity to related H1_HA proteins in NS samples from the i.m. immunized group. However, IgA binding to the seasonal-drifted H1_HA antigens was easily and uniquely detected in NS samples from the i.n. immunized animals (Figure 5I, 5M).

Given these results, we asked whether i.n. CA09HA-Ad5 immunization elicited Ab responses that could bind to more distantly related HA proteins. First, we examined whether the Abs elicited in response to CA09HA-Ad5 immunization or CA09 infection recognized H1_A/PR8/34HA (PR8/34HA) antigen, which was derived from influenza circulating in 1934 [57]. Mice immunized via the i.m. route did not make detectable PR8/34HA binding IgG in serum (Figure 5N) BAL, NW or NS (Figure S4J-L). PR8/34HA-binding IgA also could not be detected in BAL (Figure 5O), NW (Figure 5P) or NS (Figure 5Q) isolated from the i.m. immunized mice. However, mice infected or immunized via the i.n. route generated PR8/34HA-binding IgG in serum (Figure 5N), as well as PR8/34HA-binding IgA (Figure 5O-P) and IgG (Figure S4J-K) in BAL and NW. Again, only mice immunized via the i.n. route generated detectable IgA in NS that recognized the evolutionarily distant PR8/34HA antigen (Figure 5Q).

While the IgA and IgG responses elicited by i.n. immunization or infection were broadly reactive against the less conserved Group 1 H1_HA antigen, this broad responsiveness did not extend to Group 2 HA antigens, like H3_DW21HA (Figure S5A-G). However, Abs from both the i.n. immunized and CA09 infected mice exhibited detectable binding to other more distant Group 1 HA antigens, including those of avian origin like H2_NL99HA (Figure S5H-N) and H5_VN04HA (Figure S5O-U). In particular, mice immunized with CA09HA-Ad5 via the i.n. but not the i.p. route generated IgA in the LRT (BAL, Figure S5I, S5P) and the URT (NW, Figure S5J, S5Q) that bound to the H2_HA and H5_HA antigens derived from avian influenza isolates. Taken together, these data show that i.n. immunization with a replication deficient CA09HA-Ad5 vector induces a broadly reactive mucosal response that resembles the response seen after infection and is of greater magnitude and breadth than that following i.m. immunization. In addition, only i.n. immunization elicits a broadly reactive IgA response in the URT.

### Intranasal CA09HA-Ad5 immunization facilitates viral clearance and lung protection following infection with an evolutionarily distant H1 virus strain

Given that we could detect H1_PR8/34HA–reactive IgA in the URT of mice immunized i.n. but not i.m. with CA09HA-Ad5, we predicted that the i.n. immunized mice would be better protected than i.m. immunized mice following PR8/34 virus challenge. To test this, 6–8-week-old B6 mice were primed i.n. or i.m. with CA09HA-Ad5 or Empty-Ad5 and then boosted (D30) via the same route (i.n./i.n. or i.m./i.m., Figure 6A). Five days post-boost, NS were collected, and the mice were then challenged with 6500 PFU PR8/34 virus – a dose that typically causes 25-28% body weight loss in B6 mice. The infected animals were weighed daily through D15 post-challenge. On D22 post-challenge lung tissue was harvested, inflated and processed for histopathology analysis (H&E and Sirius Red staining). A separate cohort of animals was sacrificed on D3 post-challenge to measure replicating virus in the lungs and NW using the viral foci assay (VFA) (Figure 6A).

**Figure 6.**
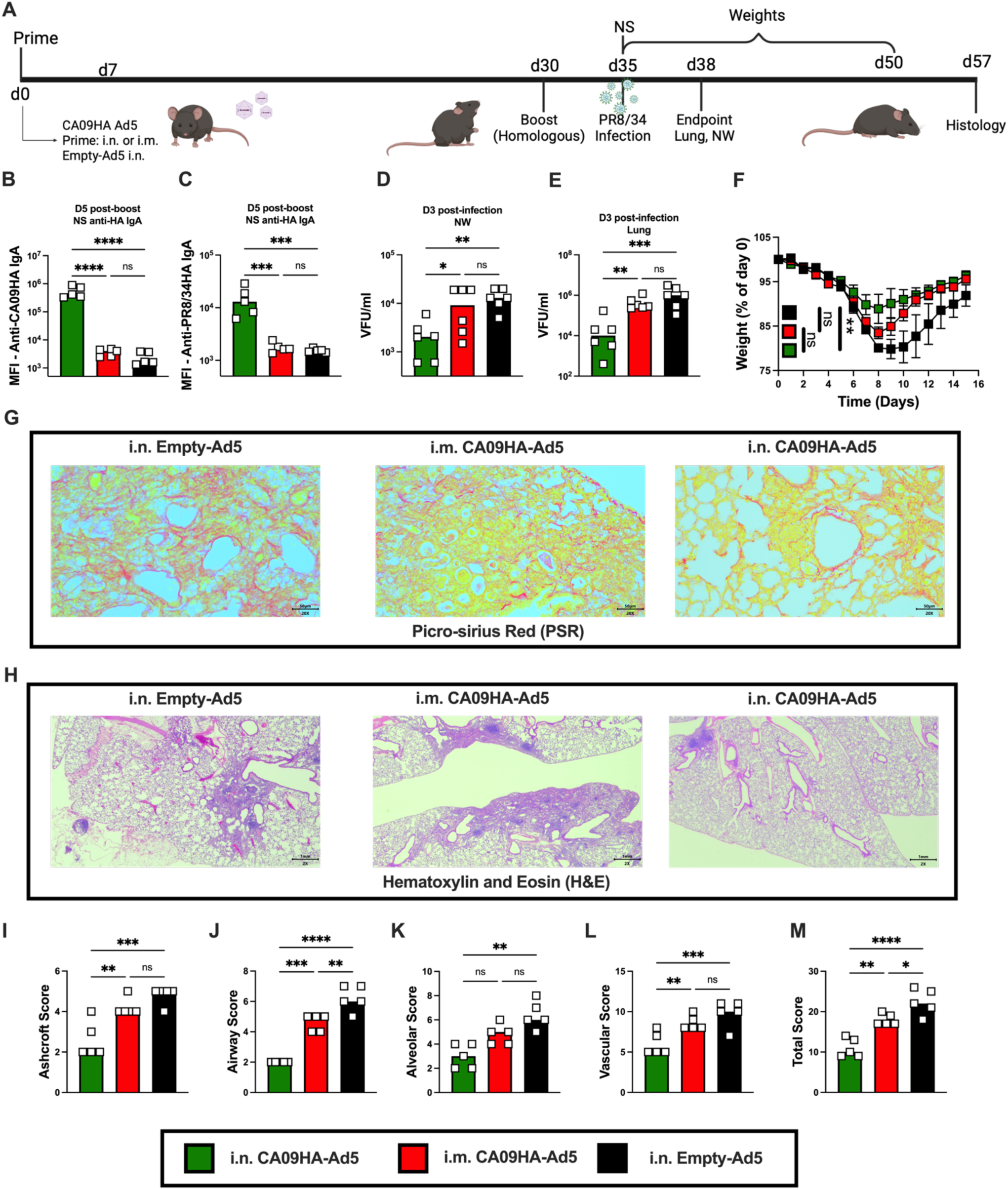
Enhanced viral clearance and decreased lung pathology in mice immunized i.n. with CA09HA-Ad5 and challenged with heterologous influenza virus. B6 mice (n=5-6/group/assay) were immunized i.n. with Empty-Ad5 (black) or CA09HA-Ad5 via the i.n. (green) or i.m. (red) route. Animals were boosted by the same route on D30 (i.n./i.n. or i.m./i.m.). NS samples were collected and analyzed by CBA D5 post-boost (n=5 animals/group). Mice were challenged i.n. with 6,500 PFU of H1_PR8/34 virus on D5 post-boost. Viral foci assays (VFA) were performed on NW and lung homogenates (n=6/group) isolated on D3 post-infection. Animals (n=5/group) were weighed daily for 15 days and lungs (n=5/group) were collected 22 days post-infection for histopathology analysis. (**A**) Study design schematic. (**B-C**) Quantitation of CA09HA-binding (**B**) and PR8/34HA-binding (**C**) IgA in NS samples collected D5 post-boost. Data reported as mean ± SD of log transformed MFI values of Ab binding to the different HA-coupled microbeads. (**D-E**) Viral burden measured by VFA in NW (**D**) and lung homogenates (**E**) D3 post-infection. Data reported as the mean ± SD of log transformed viral foci units (VFU) per ml. (**F**) Weight loss following PR8/34 infection reported as the % of starting weight and shown as mean ± SEM. (**G-H**) Representative lung tissue sections from inflated and fixed whole lungs collected on D22 post-infection. Sections stained with Picro Sirius Red (**G**) or hematoxylin and eosin (**H**). (**I-M**) Lung pathology scoring on D22 post-infection assessing lung fibrosis (Ashcroft Score, **I**), airway inflammation (**J**), alveolar involvement (**K**), vascular pathology (**L**), and total lung pathology score (**M**). See Methods for details. Scoring data reported as the mean ± SD. Data evaluated using two-way ANOVA with Tukey’s multiple comparisons test (**B-E**, **I-M**) or two-way ANOVA on AUC measurements (**F**). * p < 0.05, ** p < 0.01, *** p < 0.001, **** p < 0.0001.

Consistent with our prior data, IgA exhibiting reactivity for CA09HA antigen (Figure 6B) as well as the distantly-related PR8/34HA antigen (Figure 6C) was only detected in NS from i.n. immunized mice and was not found in NS from i.m. immunized animals. Consistent with the more broadly reactive mucosal Ab response in the i.n. immunized animals, we observed significantly less replicating PR8/34 virus in both the URT (NW, Figure 6D) and the LRT (lung, Figure 6E) of the mice immunized i.n. with CA09HA-Ad5. Indeed, at D3 post-challenge, the viral loads in NW and lungs of i.m. immunized animals were not significantly different than those observed in animals immunized with Empty-Ad5 (Figure 6D-E). Given that i.n. immunization appeared to promote more rapid clearance of the distantly related H1_PR/34 virus throughout the entire respiratory tract, it was not surprising to find that the i.n. immunized animals lost the least amount of weight and recovered more rapidly than mice immunized i.m. with CA09HA-Ad5 or i.n. with Empty-Ad5 (Figure 6F). To address whether the more rapid viral clearance seen following i.n. immunization resulted in reduced lung damage, a certified veterinary pathologist performed a blinded histopathology analysis of lung tissue sections at D22 post-challenge (Figure 6G-H). Individual scores for lung fibrosis (Ashcroft score, Figure 6I) and lung inflammation/damage in the airways (Figure 6J), alveoli (Figure 6K) and vascular (Figure 6L) compartments were determined, and an overall score (Figure 6M) was calculated (see Methods). Not surprisingly, the lungs of Empty-Ad5 immunized animals displayed marked fibrosis (Figure 6I), damage and cellular infiltration (Figure 6J-L) at D22 post-challenge. While airway infiltration was reduced in i.m. immunized animals relative to the Empty-Ad5 controls (Figure 6J), there was no difference in Ashcroft, alveolar and vascular histopathology scores (Figure 6I, 6K, 6L) between i.m. CA09HA-Ad5 immunized mice and the Empty-Ad5 immunized control mice. In fact, only the animals immunized i.n. with CA09HA-Ad5 exhibited significantly reduced lung fibrosis and inflammation (Figure 6I-L) and reduced overall pathology scores (Figure 6M). Therefore, rapid viral clearance, reduced morbidity, and prevention of lung damage following challenge with an evolutionarily distant H1 influenza virus was supported by delivery of CA09HA-Ad5 via the i.n. but not the i.m. route.

### Intranasal immunization with CA09HA-Ad5 supports broadly reactive Ab responses and accelerates viral clearance in hamsters

Intranasal immunization of laboratory mice with CA09HA-Ad5 promoted rapid viral clearance and decreased morbidity and lung damage following experimental infection with a distantly related influenza virus. While this was encouraging, it is well appreciated that human influenza viruses do not efficiently infect the URT of mice [47, 58] and that the URT of mice is anatomically distinct from humans [59, 60]. By contrast, the upper respiratory tract of Golden Syrian hamsters (GS hamsters) – a species that diverged from mice several million years ago [61, 62] – is more similar to humans [63, 64]. Moreover, GS hamsters are naturally susceptible to non-adapted human influenza A viruses, including recent H1N1 and H3N2 strains [65] and high titers of virus can be recovered from hamster nasal turbinates and washes [66, 67]. Therefore, we next asked whether i.n. delivery of CA09HA-Ad5 in hamsters also uniquely supported the development of more broadly reactive mucosal Ab responses. We immunized 9-11-week-old GS hamsters either i.n. or i.m. with 1×10⁹ VP of CA09HA-Ad5 and immunized control hamsters with Empty-Ad5 delivered via the i.n. route. All animals were given a homologous boost on D30 (i.m./i.m., i.n./i.n.) and then challenged i.n. with CA09 virus on D36. Blood and NS were collected longitudinally at multiple points and NW, and lung tissue was collected on D4 post-challenge (Figure 7A).

**Figure 7.**
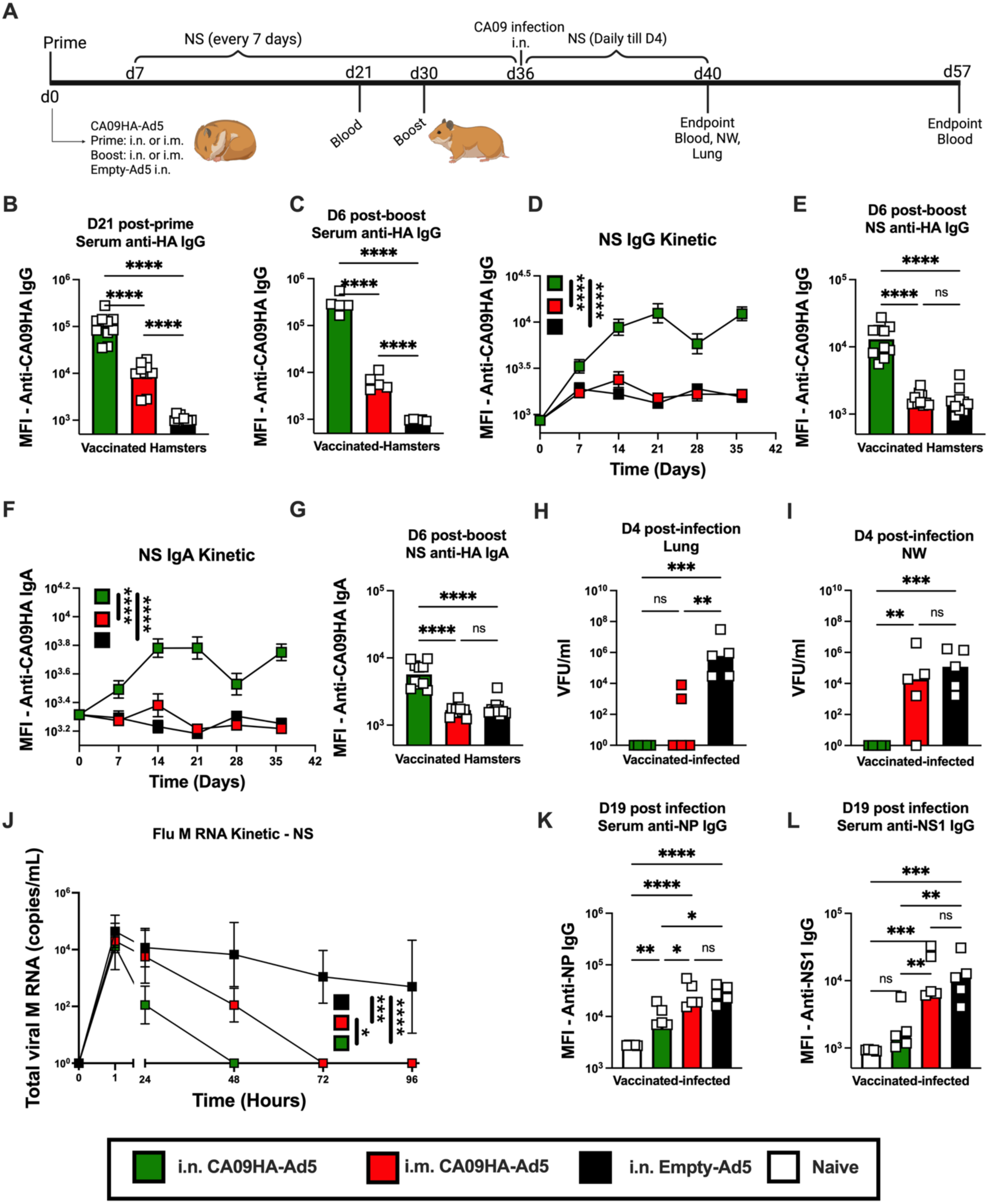
Intranasal immunization of hamsters with CA09HA-Ad5 supports broadly reactive mucosal Ab responses and accelerated viral clearance. Hamsters (n=5-10/group/assay) were immunized i.n. with Empty-Ad5 (black) or immunized with CA09HA-Ad5 via the i.n. (green) or i.m. (red) route. Animals were boosted by the same route on D30 (i.n./i.n. or i.m./i.m.) and challenged with 1×10⁸ PFU of H1_CA09 virus on D6 post-boost. NS samples were collected weekly from immunized animals (n=10/group) and for 4 days following challenge infection (n=5/group). Serum was collected from immunized animals on D21 post-prime (n=10/group), D6 post-boost (n=5/group), and D21 post-challenge infection (n=5/group). NW and lung homogenate samples (n=5/group) were collected on D4 post-challenge infection. CBA assays to measure virus-specific Abs were performed on serum and NS samples. VFA and qPCR assays to measure viral load were performed on NW, NS and lung homogenate samples. (**A**) Study design schematic. (**B-C**) Quantitation of H1_CA09HA-specific IgG in serum at D21 post-prime (**B**) and D6 post-boost (**C**). Data reported as mean ± SD of log transformed MFI values of Ab binding to CA09HA-coupled microbeads. (**D-G**) Quantitation of H1_CA09HA-specific IgG (**D-E**) and IgA (**F-G**) in NS samples collected following prime and boost immunization. Data reported as the mean ± SEM (**D, F**) or mean ± SD (**E, G**) of log transformed MFI values of Ab binding to CA09HA-coupled microbeads. (**H-I**) Viral burden measured by VFA in lung homogenate (**H**) and NW (**I**) D4 post-challenge infection. Data reported as the mean ± SD of log transformed VFU per ml. (**J**) Quantitation of viral M RNA from NS samples collected at the indicated timepoints post-infection. Data reported as the mean ± SEM of log-transformed PCR amplification data. (**K-L**) Quantitation of influenza NP-specific (**K**) and NS1-specific (**L**) IgG in serum samples collected on D21 post-infection. Data reported as mean ± SD of log transformed MFI values of Ab binding to NP-coupled or NS1-coupled microbeads. Quantitation of serum and NS IgG and IgA Ab binding to H1_BB18, H1_WI22 and H1_PR8/34 shown in Figure S6. Data analyzed using two-way ANOVA with Tukey’s multiple comparisons test (**B-C, E, G-I, K-L**) or using one-way ANOVA on AUC measurements (**D, F, J**). * p < 0.05, ** p < 0.01, *** p < 0.001, **** p < 0.0001.

Consistent with the mouse data, both i.n. and i.m. immunizations elicited CA09HA-binding IgG in serum by D21 post-prime (Figure 7B). While both routes of immunization supported the development of a systemic Ab response, i.n. immunization induced significantly higher levels of CA09HA-specific IgG in the serum compared to i.m. immunization. This remained true on D6 post-boost (Figure 7C), indicating that i.n. immunization of hamsters with a replication deficient Ad5 vector effectively supported systemic humoral immune responses. Moreover, only immunization via the i.n. route elicited CA09HA-binding IgG (Figure 7D-E) and IgA (Figure 7F-G) in NS samples. These Abs were detected as early as D7 post-primary immunization (Figure 7D, 7F) and increased following the i.n. boost (Figure 7E, 7G). Thus, i.n. immunization, but not systemic delivery of CA09HA-Ad5, induced robust URT immunity.

Next, we evaluated whether i.n. delivery of CA09HA-Ad5 selectively induced a more broadly reactive response to H1_HA antigens from recent drifted seasonal influenza A strains and historical isolates. Similar to what we previously observed in mice, IgG and IgA responses to H1_BB18HA (Figure S6A-F), H1_WI22HA (Figure S6G-L) and H1_PR8/34HA (Figure S6M-R) in serum and NS were predominantly observed in SG hamsters immunized via the i.n. route. This was particularly evident when IgA (Figure S6E-F, S6K-L, S6Q-R) and IgG (Figure S6C-D, S6I-J, S6O-P) HA-binding Abs were measured in NS samples. Moreover, and similar to what we observed in mice, boosting systemically via the i.m. route was not sufficient to initiate a local mucosal Ab response. These data therefore suggested that URT mucosal humoral immunity in hamsters also requires local Ag delivery but does not depend on the presence of replicating virus.

Given the superior mucosal immune response in the i.n. immunized SG hamsters, we predicted that these animals would be better protected following challenge infection. To test this, hamsters were challenged via the i.n. route with 1×10⁸ PFU CA09 virus at D6 post-i.n. or i.m. boost. A subset of animals was euthanized on D4 post-challenge to assess viral burden in the lungs and NW, while the remaining animals were monitored longitudinally for viral shedding in NS (Figure 7A). Consistent with robust pre-existing systemic immunity, both groups of immunized hamsters had lower levels of infectious virus in the lungs at D4 post-challenge compared to the Empty-Ad5 immunized hamsters (Figure 7H). However, while 100% of the i.n. immunized animals had no detectable virus in the lung tissue at D4 post-challenge, virus was still detected in the lung homogenates from 40% of the i.m. immunized animals (Figure 7H). Moreover, at D4 post-challenge, the viral load in NW of i.m. immunized mice was indistinguishable from the animals immunized with Empty-Ad5 (Figure 7I). By contrast, no infectious virus could be detected in NW of CA09HA-Ad5 i.n. immunized animals on D4 post-challenge (Figure 7I). Next, using a more sensitive PCR-based assay to detect influenza A matrix (M) RNA in NS, we observed equivalent levels of virus at 1h post-challenge (Figure 7J). However, by 24h post-challenge, viral M RNA levels were significantly lower in NS samples from i.n. immunized and challenged hamsters (Figure 7J) and by 48h post-challenge, no viral M RNA could be detected in NS from i.n. immunized animals (Figure 7J). Indeed, viral M RNA in nasal secretions from i.n. immunized hamsters was undetectable almost a full day earlier than i.m. immunized animals.

Since virus clearance in both the URT and LRT was significantly accelerated in the i.n. immunized hamsters, we predicted that the viral load in i.n. immunized animals might be sufficient to elicit a humoral immune response to the challenge virus. To address this, we measured the humoral immune response to virus proteins, including NP and NS1, that were not expressed by the CA09HA-Ad5 vector. Hamsters immunized with Empty-Ad5 or i.m. immunized with CA09HA-Ad5 made readily detectable serum anti-NP (Figure 7K) and anti-NS1 (Figure 7L) IgG responses, indicating that the hamsters in both groups had been productively infected with H1_CA09 virus. By contrast, i.n. immunized animals made a low to undetectable NS1 and NP IgG Ab responses following challenge (Figure 7K-L), suggesting that viral load in these animals was too low to trigger an easily measurable adaptive B cell response.

Taken together, the data show that delivery of the replication deficient Ad5-vectored antigens via the i.n. route elicited a strong systemic response and uniquely supported the establishment of URT and LRT B cells and T cell responses. Moreover, i.n. delivery of antigen using the Ad5 platform elicited more broadly reactive IgA and IgG responses within the URT, particularly at the site of viral entry. This local mucosal delivery of antigen supported accelerated viral clearance following challenge infection and resulted in less lung damage, even after challenge with a divergent strain of virus. Given that the protective response elicited by the i.n. delivered using the Ad5 platform was conserved across mice and hamsters that diverged in evolution millions of years ago [68], we argue that engagement of the mucosal immune response through local antigen delivery may be one key to developing vaccines that induce broadly reactive responses.

## DISCUSSION

The present study shows that intranasal but not intramuscular immunization with the replication defective CA09HA-Ad5 vector elicits robust mucosal B cell, T cell and IgA responses that correlate with accelerated viral clearance following challenge infection. These results, in and of themselves are not particularly surprising, and are consistent with published data comparing systemic and mucosal SARS-CoV-2 vaccination using other antigen delivery systems [44, 45]. However, we also showed that i.n. immunization of mice and hamsters with the replication defective CA09HA-Ad5 vector induces cross-reactive upper respiratory tract (URT) IgG and IgA responses. By contrast, immunization via the i.m. route that is the standard route of immunization used in humans [56], does not support establishment of the cross- or broadly-reactive IgA or IgG responses detected in either the oral pharyngeal cavity URT (captured by the NW) or the nasal secretions (captured in the nasal *swipe* (NS) samples). We further showed that i.n. immunization induces broadly-reactive IgA antibody responses in NW and NS samples – responses that were of comparable or of even great magnitude than the IgA responses elicited following natural infection. Indeed, at least in mice, i.n. immunization with CA09HA-Ad5 was superior to natural infection in eliciting a cross-reactive IgA response that could be detected in secretions found *at the surface of* the body at the interface between the nostrils and the environment. These data therefore argue that mucosal immunization with non-replicating vectors is sufficient to elicit broadly reactive antibody responses and suggest that vaccines targeting the respiratory tract, and particularly the URT, may be more effective not only in placing immune cells at the site of infection but also fundamentally changing the repertoire breadth of the responding B cells.

Although we have known for years that respiratory infection with flu is better than systemic immunization in eliciting the cross-reactive antibody response in humans [56], the reason for this dichotomy is not well-understood. An elegant prior study in mice [39] reported that influenza infection specifically supports development of lung memory B cells that express B cell receptors and antibodies that can recognize distantly related HA proteins. B cells with this broadly reactive repertoire were not present in the medLN [39], suggesting that the lung was unique in its ability to support these B cells. The lung-residing broadly-reactive memory B cells appeared to have undergone somatic hypermutation [39] and were associated with persistent germinal center responses that occur within the lung inducible bronchus associated lymphoid tissue (iBALT, [69]). The authors speculated that local ongoing GC selection in response to prolonged persistence of viral antigens within the lung and virus escape mutants might be responsible for selection of these more broadly-reactive B cells and antibodies [39]. Our data immunizing mice with a replication-deficient vector that is not targeted by the immune system and does not undergo immune mediated virus escape [49, 50, 70], suggest that the broadly reactive antibody response observed the respiratory tract is not due to selective evolution of virus in the lung or to differences in the viral antigen persistence in the lung versus other sites. Instead, the data suggest that the specialized microenvironment within the respiratory tract uniquely supports the selection of a broad more cross-reactive memory B cell and ASC repertoire.

Although replicating virus was not required to initiate a broadly reactive B cell response following i.n. immunization with CA09HA-Ad5, local antigen was clearly key to the establishment of this response in the respiratory tract. We know from work over the last decade that immune cells, including memory B and T cells, residing in the lung mucosa have distinct features compared to the recirculating populations that transit through secondary lymphoid tissues [18, 26, 31, 71]. In particular, typical germinal centers (GCs) found in tertiary lymphoid tissues like iBALT differ in important ways from the GCs present in secondary lymphoid tissues [39, 72, 73]. Indeed, the CD4 T cells that provide B cell help in the lung and support the local IgA response are phenotypically and transcriptionally distinct from conventional T_FH_ cells found in secondary lymphoid tissues [35, 74, 75]. Moreover, the follicular reticular cells and follicular dendritic cells (FDCs), which organize B cell follicles and display antigens for B cells [76], develop in response to different cues in the lung (inflammatory signals) and LNs (embryonic programming). Given that the combination of antigen and T cell help controls B cell selection within the GC [77], it is possible that the lung GC B cell response induced in response to flu infection or immunization with CA09HA-Ad5 is less restrictive and permits the survival of local memory B cells and ASCs that, while mutated, are not highly selected and continue to retain a broad reactivity profile.

It is also possible that the memory B cells generated in secondary lymphoid tissues and then take up residence in the lung [31, 78] are different from those memory B cells that remain in circulation. We know that lung BRM cells can arise from the systemic LN response [34] but once in the lung exhibit distinct transcriptional programs relative to memory cells from secondary lymphoid tissues [79–82]. It is postulated that these tissue residency transcriptional programs support maintenance of cells in the mucosa and likely contribute to the functional properties of these local cells. Indeed, data from our group and others show that development of the non-recirculating resident flu-specific memory B cell compartment requires the presence of antigen in the lung [23, 34] as well as IFNγ-producing T cells [37]. We further reported [38] that flu-specific B cells receiving IFNγ-dependent signals from T cells upregulate expression of the transcription factors STAT1 and T-bet and take on an “effector-like” memory transcriptional profile. We found that maintenance of the lung effector memory B cell compartment required continuous expression of the IFNγ-induced transcription factor T-bet and that sustained expression of T-bet by the lung effector memory B cells was necessary for the rapidly differentiation of these cells into ASCs in the lung following reactivation [38]. Thus, local antigen as well as cytokine signals appear to be important for establishment of the broadly reactive memory B cell subset and the lung-resident memory B cell compartment.

Our new data do not discriminate whether all lung-resident memory B cells are also broadly reactive. However, our data show that local infection with a replication deficient virus is sufficient to establish the T-bet expressing memory B cell compartment and the URT broadly reactive antibody response. Although we know that these lung resident effector T-bet^+^ B cells include the memory B cells with rapid differentiation potential following reactivation [38], these cells primarily differentiate into IgG-producing ASCs following challenge infection [38]. Thus, while the lung resident T-bet^+^ memory B cells may be responsible for the enhanced cross-reactive IgG responses detected in the BAL (LRT) and NW (URT) of the i.n. immunized mice, it is unlikely that the T-bet^+^ memory B cells in the lung and BAL are solely responsible for producing the cross-reactive IgA antibodies that we detect only after i.n. immunization in the NS samples that catch nasal secretions at the body surface. Instead, we postulate that the memory B cells that give rise to the broadly reactive IgA^+^ antibody may be localized within the nasal mucosa akin to those recently described in human nasal mucosa [81].

Regardless which B cell population(s) in the respiratory tract contribute to the broadly reactive B cell and antibody repertoire within this site, our findings show that i.n. administration of CA09HA-Ad5 to both mice and hamsters, which diverged in evolution millions of years ago [68], was sufficient to induce flu HA-specific IgG and IgA antibodies that could bind to antigenically drifted H1 isolates (H1_BB2018, H1_HI2019) and even the distally related H1_PR8/34 strain that circulated almost 100 years ago. Moreover, only i.n. administration of CA09HA-Ad5 induced IgA and IgG capable of binding to H2 and H5 avian HA antigens in the BAL. Similarly, only i.n. immunization with CA09HA-Ad5 elicited IgA antibodies in the BAL and NW that could bind the avian HA antigens. While we cannot say with certainty that the increased URT protection observed in the i.n. immunized animals that were subsequently challenged with heterologous virus was solely due to the presence of cross-reactive antibodies in the URT, we postulate that the presence of these non-neutralizing but broadly reactive antibodies could support antigen presentation, enhance NK cell activation and facilitate rapid antigen clearance [83]. This more rapid clearance was also associated with minimal lung tissue damage as animals immunized via the i.n. route exhibited significantly reduced inflammation, fibrosis, and tissue remodeling compared to animals immunized i.m. or with the empty vector control. Given that severe influenza is often associated with chronic lung pathology and impaired respiratory function [84–87], these findings highlight the clinical relevance of mucosal immunization in preserving lung health after infection.

Taken together, our data suggest that i.n. immunization with an Ad5-vectored HA construct elicits superior protection compared to traditional i.m. delivery. Indeed, i.n. immunization with CA09HA-Ad5 uniquely established local lung and lung airway memory B and T cell responses and facilitated the URT IgA response. Moreover, i.n. immunization with this replication-deficient Ad5 vector was sufficient to induce broadly reactive IgA responses directly at the site of virus entry. Secretory IgA has long been recognized as a critical barrier for respiratory pathogens, providing both neutralization at the site of viral entry and immune exclusion that limits viral spread across mucosal barriers [21, 22, 88–92]. These URT IgA Abs, which were exclusively induced following immunization via the i.n. route, were broadly reactive and could bind HA antigens from seasonal drift viral variants, historic virus strains and even HA antigens from avian influenza viruses. Given the long-term goal of developing a universal influenza vaccine [93–96], it seems reasonable to conclude that one important component will be to deploy vaccine vectors, like the Ad5 [E1-, E2b-E3-] platform, that can be administered via the mucosal route and will elicit local respiratory tract immune responses including the broadly reactive IgA in the URT that can support rapid clearance viral variants directly at the site of infection.

## Author Contributions

F.E.L., T.D.R., M.D.S., A.S-S., and D.B. conceived and conceptualized the study. F.E.L and T.D.R. secured funding. F.E.L., T.D.R. and D.B. provided direct supervision of the project. F.E.L., M.D.S., A.S-S., and D.B. designed the experiments that were performed by M.D.S., D.B., A.S-S., I.R.C., D.K., S.M.T., J.A.H., D.M.F. and F.D. E.S.G. and A.E.R. developed, produced and provided the Ad5 vectors for the experiments in this manuscript. G.Y., J.A. and K.S.H. prepared immunoassay reagents. A.C.K.L., F.Z., J.A., M.E.C. and W.O.S. performed the immunoassays. D.Y. performed the viral RNA quantitation with oversight from S.M.L. Board-certified veterinary pathologist J.B.F. performed the histopathology analyses. M.D.S. and F.E.L. curated the data. M.D.S. and F.E.L wrote the manuscript and prepared the final figures. All authors had full access to the data in the study and reviewed the manuscript.

## Acknowledgements

The authors would like to thank Thomas ‘Scott’ Simpler, Uma Mudunuru, Rebecca Burnham and Kelsey Browning (UAB) for animal husbandry; Derek B. Moates and Kamaljeet Kaur (UAB Fungal Reference Lab) for coordinating sample drop-off and data delivery; Emily E. Helmen and Sherri Coffman (UAB Comparative Pathology Lab) for histology; Dr. Andrea J. Osborne, DVM (UAB Animal Resources Program) for veterinary care of animals; Dr. Douglas M. Fox (SEBLAB Research Operations Manager), Dr. Francisco Dominguez, Dr. Joseph W. Palmer (SEBLAB Research Safety Manager) for facilitating work in the SEBLAB ABSL2 space. Funding for experiments was provided by the NIH: R01 AI110508 (to F.E.L.), R01 AI150740 (to F.E.L.), R01 AI153413 (to T.D.R.) and R01 AI52476 (to T.D.R.). NIH P30 CA013148 and P30 AI027767 provided support for the UAB consolidated flow cytometry core. NIAID Regional Biocontainment Lab (RBL) awards UC6AI058599, G20AI167409, and UC7AI180255 provided support for the UAB SEBLAB Regional Biocontainment Laboratory. We thank the NIH Tetramer Core Facility (NIH Contract 75N93020D00005 and RRID:SCR_026557) for providing the H2-Db Influenza A NP 366-374 tetramer that was used in this study, and the UAB Immunology Institute Antibody Characterization and Serology facility for virus-specific antibody CBAs. Infographics were generated using Biorender.

## Declaration of Interests

E.S.G. and A.E.R. are employed at ImmunityBio, which provided the Ad5-based vectors used in this study. The other authors declare no competing interests.

## Methods

### Animal studies

Nine-week-old male Lakeview Golden Syrian hamsters (GS hamsters) were obtained from Charles River Laboratories and were housed within the University of Alabama at Birmingham (UAB) animal facilities, including the Southeastern Biosafety Laboratory (SEBLAB) ABSL-3 facilities. C57BL/6 (B6) mice, which were obtained from Jackson laboratories, and B6.T-bet-ZsGreen T-bet reporter mice [97] which were acquired from Dr. Zhu (NIH), were bred in the UAB animal facilities under specific pathogen-free conditions. All procedures were approved by the UAB Institutional Animal Care and Use Committee (IACUC protocol 22065) and the UAB Biosafety Committee (protocols 20-083) and were conducted in accordance with guidelines from the National Research Council.

### Production of Adenovirus vectors

The replication deficient adenovirus delivery platform Ad5 [E1-, E2b-, E3-] [49], which contains deletions in the early 1 (E1), early 2 (E2b) and early 3 (E3) regions, was constructed as previously described [48, 70]). The parent Ad5 [E1-, E2b-, E3-] vector (Empty-Ad5 or Ad5-null) was modified [51] to express CA09HA (Ad5 [E1-, E2b-, E3-] CA09HA, abbreviated as CA09HA-Ad5) or influenza NP (Ad5 [E1-, E2b-, E3-] NP, abbreviated as NP-Ad5). The 1743 base pair coding sequence for the CA09HA insert represents the consensus gene sequence generated from the pandemic 2009 H1N1 strains that circulated in North and South America. The 1565 base pair coding sequence for the NP insert was derived from the GD96 H5N6 avian influenza A strain (https://www.ncbi.nlm.nih.gov/nuccore/nc_007360). Empty-Ad5, CA09HA-Ad5 and NP-Ad5 vectors were produced as previously described and stocks of virus at 1 x 10^12^ viral particles (VP)/mL were frozen at -80°C until used [48].

### Immunizations

Seven-week-old mice and twelve-week-old male GS hamsters were immunized with Empty-Ad5, CA09HA-Ad5 or NP-Ad5, administered via the intranasal (i.n.), intraperitoneal (i.p.), or intramuscular (i.m.) route. 1 × 10⁹ VP of Empty-Ad5, CA09HA-Ad5 or NP-Ad5 was delivered in a total volume of 100 µL via the i.n., i.p., and i.m. route. In some experiments, animals were given a second equivalent booster dose of the same Ad5 vector delivered by the same route (homologous boost) 30 days following the prime dose. All animals were sedated with isoflurane during i.n. and i.m. administration and inoculated in the supine position.

### Influenza infections

Animals were anesthetized with isoflurane and inoculated intranasally (i.n.) with the indicated influenza A virus strains diluted in sterile phosphate-buffered saline (PBS). Mice were infected with either H1N1 A/Puerto Rico/8/1934 (PR8/34) at a sublethal dose of 6.5 × 10³ VFU in a 100 µL inoculum, or with the H1N1 A/California/04/2009 (CA09) strain at a dose of 5.0 × 10² VFU in the same volume. GS hamsters were infected i.n. with 1 × 10⁸ VFU CA09 diluted in PBS (total volume 100 µL). Following infection, animals were monitored daily for body weight, clinical signs, and survival for the duration of the study.

### Nasal swipe collection

Nasal swipes (NS), which are used to sample Ab and virus present at the interface between the nasal cavity and the outside environment, were collected from hamsters and mice. To limit movement of the animals, mice were gently restrained, and hamsters were placed in an open but constrained container. The external surface of the hamster or mouse nose was swiped for 5-10 seconds (sec) using a polyester-tipped swab (Medical Packaging SP-7D) that was pre-moistened in a screw cap tube containing 300 µL of viral transport medium (VTM; HBSS containing Ca²⁺ and Mg²⁺, supplemented with 2% FBS, 100 µg/mL gentamicin, and 0.5 µg/mL amphotericin B). The swab tip was cut, and the tip was placed back in the same VTM-containing screw-cap tube. The tubes were vortexed briefly, and swab tips were discarded. Samples were aliquoted and stored at -80°C until analyzed using a cytometric bead array (CBA) or Viral Foci Assay (VFA).

### Nasal wash collection

Nasal wash (NW), which captures virus and Abs present in the oral pharyngeal and nasal cavities of the upper respiratory tract (URT) [98], were collected post-mortem at the indicated timepoints. Animals were euthanized by i.p. injection of tribromoethanol (3 mL/hamster or 600 µL/mouse). The trachea was surgically exposed, and a small incision was made to allow insertion of a 19-gauge blunt needle attached to a 1-mL insulin syringe. VTM (400 µL) was flushed through the nasal passages via the trachea while the mouth was held closed, allowing fluid to exit through the nostrils and be collected into a screw-cap tube. Samples were aliquoted and stored at -80°C until analyzed by CBA or VFA.

### BAL collection

Bronchoalveolar lavage fluid (BAL), which captures virus and Abs present in the lung airway and lower respiratory tract (LRT), was collected postmortem from mice by inserting ethyl vinyl acetate (EVA) microbore tubing (0.02 in ID, 0.06 in OD; Cole-Parmer) into the exposed trachea. The tubing was connected to a 23G needle and 3-way stopcock attached to a syringe. A single 1 mL lavage of the lungs was performed using ice-cold, VTM, supplemented with 2 mM EDTA. Recovered fluid was centrifuged at 1,800 rpm for 5 minutes (min) to separate cells from the supernatant. Samples were aliquoted and stored at -80°C until analyzed by CBA or VFA.

### Serum collection

Blood samples were collected from sedated hamsters via the saphenous vein at multiple time points post-immunization. Terminal blood samples were collected following infection with CA09. In mice, blood was collected at the time of euthanasia. All samples were collected into BD Microtainer® blood collection tubes (BD Biosciences) and centrifuged at 10,000 rpm for 10 min at room temperature (RT) to separate serum. Serum was aliquoted and stored at -80°C until analyzed.

### Lung processing for viral foci assays (VFA)

Lung tissues were harvested from hamsters and mice following euthanasia. For hamster samples, the left and right lung lobes were dissected and placed into separate 2 mL Lysing Matrix M tubes (MP Biomedicals, #116923050-CF), each containing 1 mL VTM. Mouse lungs were processed whole, and the entire lung was placed into a single Lysing Matrix M tube containing 1 mL VTM. Tissues were homogenized using a FastPrep-24 Classic bead-beating grinder lysis system (MP Biomedicals, #116004500), set to 4.0 m/s for 20 seconds. Homogenized samples were placed on ice for 5 min, then subjected to a second homogenization using the same settings. After processing, samples were centrifuged at 1800 rpm for 5 min at RT to pellet debris. For hamster samples, 300 μL of supernatant was collected from each homogenized lobe and combined to generate a representative whole-lung sample. Samples were aliquoted and stored at -80°C until analyzed by VFA.

### VFA protocol

To measure replication competent influenza virus in mouse and hamster samples, VFA were performed as previously described [99]. Briefly, MDCK cells were seeded at 5 × 10³ cells per well in 96-well tissue culture-treated plates (200 µL/well of MEM supplemented with 10% FBS, 2% Penicillin-Streptomycin/L-glutamine, and 1% Amphotericin B) and incubated at 37°C, 5% CO₂ for 4 days. On D5, experimental samples were serially diluted (10-fold dilutions) in 96-well V-bottom plates containing virus diluent (MEM + 4 µg/mL trypsin). The MDCK monolayers were washed twice with HBSS, and 100 µL of each dilution was transferred to the MDCK containing wells. Plates were centrifuged at 2,000 rpm for 1 hr at RT, washed twice with virus diluent, and incubated overnight at 33°C with 5% CO₂. On day 6, the MDCK monolayers were fixed with cold 80% acetone in PBS (-20°C, 20 min). Cells infected with influenza A virus were enumerated using a mouse anti-influenza A-biotin primary antibody (Millipore, lot no. 4083751) followed by streptavidin-alkaline phosphatase secondary antibody (SouthernBiotech, Catalog no. 7105-04). Viral foci were detected with BCIP/NBT substrate and counted using a dissecting microscope and multiplied by inoculum and viral dilution to determine VFU/ml.

### Histopathology

Mouse lungs were inflated with 1 mL 10% neutral-buffered formalin (NBF) using a blunt 19-gauge needle attached to a 5-mL syringe, then fixed for 7 days in 50-mL conical tubes containing 10% NBF. Fixed tissues were embedded dorsal side down in paraffin, sectioned at 5 µm, and slides containing lung sections were prepared. Lung section slides were stained with hematoxylin and eosin (H&E) or Picro Sirius Red (PSR). Representative images were captured using a Nikon Eclipse Ci microscope and analyzed with NIS-Elements software (Nikon). Histopathological scoring was performed by a board-certified veterinary pathologist, who was blinded to experimental groups. As previously described [98], airway, alveolar, vascular involvement and injury were scored with the extent of involvement (scored from 0-4) and severity as it pertains to inflammation, injury, and reactive epithelial/endothelial changes (scored from 0-4) for each compartment summed individually. To determine the extent of fibrosis within the alveolar region (scored from 0-8), PSR positive staining was evaluated at 20x magnification to generate the Ashcroft Score [100]. This score was added to vascular involvement and injury measurements obtain the total vascular score. Airway, alveolar, vascular, and fibrotic pathology were each scored in affected regions, and the scores were summed to generate a total lung pathology score. See Table S1 for score rubric.

### Quantitative PCR measurement of viral M RNA

Nasal swipe samples (300 µL) were collected from individual hamsters and mixed 1:1 with lysis buffer containing 10% proteinase K from the Maxwell® RSC Viral Total Nucleic Acid Purification Kit (Promega, #AS1330). Samples were vortexed and incubated at 56°C for 10 min for viral inactivation prior to RNA extraction using the Maxwell® RSC 48 Instrument (Promega, Madison, WI). Quantification of influenza A viral N RNA was performed via qRT-PCR on a QuantStudio™ 5 Real-Time PCR System (ThermoFisher, Waltham, MA) using the following universal influenza A virus primers that target the matrix (M) gene [101]: Forward primer 5′-CAA GAC CAA TYC TGT CAC CTY TGA C-3′, Reverse primer 5′-GCA TTT TGG ATA AAG CGT CTA CG-3′ and Probe 5’-/FAM/TGC AGT CCT /Nova/ CGC TCA CTG GGC ACG/BHQ-1/-3’. No template PCR and no template extraction controls were incorporated into each run. Thermocycling parameters included: RT incubation at 50°C for 5 min, Taq activation at 95°C for 20 sec, 45 cycles at 95°C for 3 sec and 55°C for 30 sec.

### Production of recombinant influenza HA, NP and NS1 proteins

Recombinant influenza HA, NP and NS1 proteins were produced as previously described [33, 102, 103]. Briefly, synthetic constructs (GeneArt, Regensburg Germany) containing influenza HA ectodomains were cloned into the pCXpoly(+) vector modified with a human CD5 leader sequence and a GCN4 isoleucine zipper trimerization domain (GeneArt) followed by either a 6X-His tag (HA-6XHIS construct) or an AviTag (HA-AviTag construct). The HA–6XHIS and HA–AviTag constructs for each HA were co–transfected using 293Fectin Transfection Reagent into FreeStyle™ 293–F Cells (ThermoFisher Scientific) at a 2:1 ratio [102].Transfected cells were cultured in FreeStyle 293 expression medium (ThermoFisher Scientific) for 3 days. Culture supernatant was collected, and the recombinant HA trimeric proteins were purified by FPLC using a HisTrap HP Column (GE Healthcare). Trimeric HA proteins were biotinylated *in vitro* using BirA enzyme (Avidity), then buffer exchanged and concentrated. The coding sequence of NP from the PR8/34 virus was synthesized (GeneArt) in frame with the coding sequence for a 15 amino acid biotinylation consensus site [104] added to the 3′ end. The modified NP sequence was cloned in frame with a 6X-HisTag in the pTRC–His2c expression vector (Invitrogen). NP protein was expressed in *E. coli* strain CVB101 (Avidity) that co-expresses the BirA biotin ligase. NP protein was purified by FPLC using a HisTrap HP Column and eluted with a 50–250 mM imidazole gradient. The eluted protein was buffer exchanged and concentrated. The PR8/34 NS1 gene was chemically synthesized (GeneArt) in frame with a 3’ addition of the coding sequence for the BirA enzymatic biotinylation consensus site and the 6X-HisTag. Two mutations, R38A and K41A, demonstrated to prevent aggregation of NS1 at high concentrations [105], were introduced into the NS1 gene by site-directed mutagenesis. The recombinant NS1 gene was cloned into the pTRC-His2c expression vector and expressed in the BirA-enzyme containing *E. coli* strain CVB101. Biotinylated recombinant NS1 was purified by FPLC using a HisTrap HP Column, eluted with a 50-250 mM imidazole gradient, buffer exchanged and concentrated. Biotinylated recombinant HA_CA09 trimers and NP monomers were tetramerized with fluorochrome-labeled streptavidin (Agilent), centrifuged as 20,000 x g for 10 min at 4°C and then aliquot and store in the dark at 4°C. See Table S2 for description of all flu antigens assessed in this study.

### Preparation of lung, MedLN, and BAL single cell suspensions

Lung, MedLN, and BAL samples were collected from mice at experimental endpoints. Lungs were mechanically dissociated and incubated at 37°C for 30 min in PBS containing collagenase (1.25 mg/mL; Millipore-Sigma, Burlington, MA, USA) and DNase I (150 U/mL; Millipore-Sigma). Following digestion, lung tissue was washed and passed through a 100μm cell strainer to prepare single-cell suspensions. Red blood cell (RBC) lysis was performed using ACK lysis buffer, and samples were filtered again to remove residual debris. MedLN tissues were mechanically dissociated, washed and filtered. BAL fluid was centrifuged to isolate cells, which were resuspended in PBS, treated with RBC lysis buffer, and filtered through a 100μm strainer to obtain single-cell suspensions. All cell suspensions were kept on ice or at 4°C until further processed.

### Cell recovery

Cell recovery from samples was determined by plating 20 μL of the single-cell suspensions into 96-well V-bottom plates and staining with the live/dead dye 7-Aminoactinomycin D (7-AAD; BioLegend, Catalog no. 420403) at 4°C for 15 min. Cells were washed and resuspended in 50 μL of staining media containing Fluoresbrite® Carboxylate YG 10 μm microspheres (2 × 10⁵ beads/mL; Polysciences Inc., Warrington, PA, USA). Samples were analyzed by flow cytometry (see below) and the number of beads and live cells in each sample was used to calculate the number of total live cells present in each sample.

### Flow cytometry phenotyping

200 μL of each single cell suspension was transferred to 96-well V-bottom plates and centrifuged at 1800 rpm for 3 min. Cells were then resuspended in 50 μL of staining buffer (PBS, and 2% FBS) and Abs at the indicated concentration (See Table S3 for Abs used in this study). Cells were incubated at 4°C for 20 min in the dark. Following surface staining, cells were washed once with staining buffer, then incubated with a LIVE/DEAD viability dye. After viability staining, cells were washed and resuspended in 200 µL of staining buffer for flow cytometric analysis. Data was acquired on either a BD FACSCanto II or a BD FACSymphony flow cytometer (BD Biosciences, East Rutherford, NJ, USA). Data were analyzed using FlowJo software v10.10.0 (BD Biosciences), with standard gating strategies based on forward and side scatter, singlet discrimination, and fluorescence minus one (FMO) controls where applicable.

B cell subsets were identified using CD19 (Biolegend, Catalog no. 115558), CD138 (Biolegend, Catalog no. 142508), IgD (Biolegend, Catalog no. 405723), CD38 (eBioscience, Catalog no. 25-0381-82), CD95 (BD, Catalog no. 554257), and PNA (Life Tech, Catalog no. L21409). Influenza-specific B cells were identified using CA09HA trimers that were tetramerized and conjugated to SA-PE and SA-APC or using NP monomers that were tetramerized and conjugated to SA-APC (see above). Abs used to exclude macrophages and T cells (Dump Abs) included CD64 (BioLegend, Catalog no. 139308) and CD3ε (Invitrogen, Catalog no. 45-0031-82). 7AAD (BioLegend, Catalog no. 420403) was used to exclude dead cells. Zs-Green was used to identify T-bet expressing B cells in the T-bet reporter mice.

T cell subsets were identified using CD3 (BioLegend, Catalog no. 155612), CD4 (BD, Catalog no. 560782), and CD8α (BD, Catalog no. 561967). Dump channel antibodies used to exclude macrophages and monocytes included CD64 (BioLegend, Catalog no. 139308) and CD11b (BD Pharmingen, Catalog no. 550993). Live/Dead Aqua viability dye (ThermoFisher, Catalog no. L34965) was used to exclude non-viable cells. Influenza NP-specific CD8⁺ T cells were identified using H-2D(b) NP₃₆₆-₃₇₄ tetramers (NIH Tetramer Core Facility, Emory University, Lot no. 78600 and 77852).

A second flow cytometry panel was prepared to identify tissue-resident memory (TRM), effector (Teff), and memory (Tmem) T cell populations. Dead cells were excluded with Live/Dead Aqua, and B cells were excluded using CD19 (BioLegend, Catalog no. 115543). CD8⁺ T cells were identified using CD8 (BD, Catalog no. 612898) in combination with CD62L (BD, Catalog no. 560514) and CD3ε (BD, Catalog no. 750638). NP-specific CD8⁺ T cells were identified using CD44 (eBioscience, Catalog no. 12-04441-83) and the H-2D(b) NP_366-374_ tetramer. TRM cells were defined as CD69⁺CD103⁺ (BioLegend, Catalog no. 104529; BD Horizon, Catalog no. 564320, respectively). Teff and Tmem populations were characterized using CXCR3 (BD, Catalog no. 751457) and CXCR6 (BioLegend, Catalog no. 151109) expression patterns.

### Generation of influenza-specific CBA reagents

Generation of HA-specific, NS1-specific and NP-specific CBA reagents has been previously described [102]. Briefly, biotinylated recombinant influenza NP, NS1, or HA proteins (See Table S2) were coupled to streptavidin functionalized 4-μm or 5-μm fluorescent microparticles (Carboxyl Blue Particle Array Kit, Spherotech Inc.). Each biotinylated recombinant protein, 500 μg, was incubated with 2 x 10^7^ streptavidin functionalized fluorescent microparticles in 400 μL of 1% BSA in PBS. Protein-coupled beads were washed and resuspended at 1 × 10⁸ beads/mL and stored at 4°C until used.

### CBA measurement of influenza-specific IgG and IgA responses

Forty μL aliquots of serum (diluted 1:2,000 in PBS), NW, or NS samples were arrayed in 96-well U-bottom polystyrene plates. A suspension containing 14 distinct influenza A antigen-coated bead sets (3 x 10^3^ cytometric beads per set) was added in 5 μL volume to each sample. Suspensions were gently mixed, incubated for 15 min at RT, washed in PBS, then stained for 15 min at RT with species-specific fluorescein isothiocyanate (FITC)-conjugated polyclonal IgG or phycoerythrin (PE)-conjugated polyclonal anti-IgA (SouthernBiotech). After staining, the antigen-bound beads were washed with PBS and resuspended in 100 μL PBS. Samples were collected on a Beckman Coulter CytoFLEX flow cytometer in plate mode at a sample rate of 100 μL/min for 1 min. After acquisition, the FCS files were analyzed using FlowJo software (BD Biosciences). The beads were identified by gating on singlet 4-μm or 5-μm particles using forward and side scatter in log scale. Distinct bead populations, each corresponding to a specific influenza A antigen, were identified based on APC-Cy7 channel fluorescence. Geometric mean fluorescence intensity (gMFI) was calculated in the FITC or PE channel.

### Quantification and statistical analysis

Statistical analysis of all experiments including tests used, n, and number of experimental repeats are provided in figure legends. FlowJo (v10, Tree Star) was used for flow cytometric analyses. GraphPad Prism (v10.3.0) was used for statistical analyses and data visualization.

### Lead Contact

Further information and requests for recombinant flu reagents should be directed to the lead contact, Frances Lund.

### Materials Availability

Recombinant influenza NP, HA and NS1 expression constructs can be provided by the lead contact. Recombinant proteins used in this paper and fluorochrome-labeled HA- and NP-tetramer reagents can be ordered through the UAB Immunology Institute Antibody Characterization and Serology Core <u>(UAB II</u> <u>ACS).</u> Breeder pairs for mouse strains are available upon request but require MTA and regulatory approval before shipment. To enquire about the Ad5 constructs used in this manuscript, contact E.S.G. at ImmunityBio.

### Data and code availability

All data reported in this paper will be shared by the lead contact upon request. No original code was reported in this manuscript.

## Notes

### Competing Interest Statement

E.S.G. and A.E.R. are employed at ImmunityBio, which provided the adenovirus serotype 5-based vectors used in this study. The other authors declare no competing interests.

## References

1. Molinari, N.A., et al., The annual impact of seasonal influenza in the US: measuring disease burden and costs. Vaccine, 2007. 25(27): p. 5086–96.

2. Fireman, B., et al., Influenza Vaccination and Mortality: Differentiating Vaccine Effects From Bias. American Journal of Epidemiology, 2009. 170(5): p. 650–656.

3. Paget, J., et al., Global mortality associated with seasonal influenza epidemics: New burden estimates and predictors from the GLaMOR Project. J Glob Health, 2019. 9(2): p. 020421.

4. Macias, A.E., et al., The disease burden of influenza beyond respiratory illness. Vaccine, 2021. 39: p. A6–A14.

5. Chen, J.R., et al., Better influenza vaccines: an industry perspective. Journal of Biomedical Science, 2020. 27(1).

6. Gouma, S., E.M. Anderson, and S.E. Hensley, Challenges of Making Effective Influenza Vaccines. Annual Review of Virology, Vol 7, 2020, 2020. 7: p. 495–512.

7. Becker, T., et al., Influenza Vaccines: Successes and Continuing Challenges. J Infect Dis, 2021. 224(12 Suppl 2): p. S405–S419.

8. Kimball, J., et al., Influenza Vaccine Failure Associated With Age and Immunosuppression. Journal of Infectious Diseases, 2021. 224(2): p. 288–293.

9. White, E.B., et al., Influenza Vaccine Effectiveness Against Illness and Asymptomatic Infection in 2022-2023: A Prospective Cohort Study. Clin Infect Dis, 2025. 80(4): p. 893–900.

10. Li, G.M., et al., Pandemic H1N1 influenza vaccine induces a recall response in humans that favors broadly cross-reactive memory B cells. Proc Natl Acad Sci U S A, 2012. 109(23): p. 9047–52.

11. Margine, I., et al., H3N2 influenza virus infection induces broadly reactive hemagglutinin stalk antibodies in humans and mice. J Virol, 2013. 87(8): p. 4728–37.

12. Moody, M.A., et al., H3N2 influenza infection elicits more cross-reactive and less clonally expanded anti-hemagglutinin antibodies than influenza vaccination. PLoS One, 2011. 6(10): p. e25797.

13. Pica, N., et al., Hemagglutinin stalk antibodies elicited by the 2009 pandemic influenza virus as a mechanism for the extinction of seasonal H1N1 viruses. Proc Natl Acad Sci U S A, 2012. 109(7): p. 2573–8.

14. Wrammert, J., et al., Broadly cross-reactive antibodies dominate the human B cell response against 2009 pandemic H1N1 influenza virus infection. J Exp Med, 2011. 208(1): p. 181–93.

15. Krammer, F., The human antibody response to influenza A virus infection and vaccination. Nat Rev Immunol, 2019. 19(6): p. 383–397.

16. Reperant, L.A., G.F. Rimmelzwaan, and A.D. Osterhaus, Advances in influenza vaccination. F1000Prime Rep, 2014. 6: p. 47.

17. McMillan, C.L.D., et al., The Next Generation of Influenza Vaccines: Towards a Universal Solution. Vaccines (Basel), 2021. 9(1).

18. Mettelman, R.C., E.K. Allen, and P.G. Thomas, Mucosal immune responses to infection and vaccination in the respiratory tract. Immunity, 2022. 55(5): p. 749–780.

19. Brandtzaeg, P., Role of mucosal immunity in influenza. Dev Biol (Basel), 2003. 115: p. 39–48.

20. Wang, T., F. Wei, and J. Liu, Emerging Role of Mucosal Vaccine in Preventing Infection with Avian Influenza A Viruses. Viruses, 2020. 12(8).

21. de Fays, C., et al., Secretory Immunoglobulin A Immunity in Chronic Obstructive Respiratory Diseases. Cells, 2022. 11(8).

22. Sinha, D., et al., Unmasking the potential of secretory IgA and its pivotal role in protection from respiratory viruses. Antiviral Res, 2024. 223: p. 105823.

23. Oh, J.E., et al., Intranasal priming induces local lung-resident B cell populations that secrete protective mucosal antiviral IgA. Sci Immunol, 2021. 6(66): p. eabj5129.

24. Booth, J.S. and F.R. Toapanta, B and T Cell Immunity in Tissues and Across the Ages. Vaccines (Basel), 2021. 9(1).

25. Nelson, S.A. and A.J. Sant, Potentiating Lung Mucosal Immunity Through Intranasal Vaccination. Frontiers in Immunology, 2021. 12.

26. Humphries, D.C., et al., Pulmonary-Resident Memory Lymphocytes: Pivotal Orchestrators of Local Immunity Against Respiratory Infections. Front Immunol, 2021. 12: p. 738955.

27. Alqahtani, S.A.M., Mucosal immunity in COVID-19: a comprehensive review. Front Immunol, 2024. 15: p. 1433452.

28. Davis-Porada, J., et al., Maintenance and functional regulation of immune memory to COVID-19 vaccines in tissues. Immunity, 2024. 57(12): p. 2895–2913.e8.

29. Mitsi, E., et al., Respiratory mucosal immune memory to SARS-CoV-2 after infection and vaccination. Nat Commun, 2023. 14(1): p. 6815.

30. Terreri, S., et al., Persistent B cell memory after SARS-CoV-2 vaccination is functional during breakthrough infections. Cell Host Microbe, 2022. 30(3): p. 400–408 e4.

31. Chen, C. and B.J. Laidlaw, Development and function of tissue-resident memory B cells. Adv Immunol, 2022. 155: p. 1–38.

32. Longet, S. and S. Paul, Pivotal role of tissue-resident memory lymphocytes in the control of mucosal infections: can mucosal vaccination induce protective tissue-resident memory T and B cells? Front Immunol, 2023. 14: p. 1216402.

33. Allie, S.R., et al., The establishment of resident memory B cells in the lung requires local antigen encounter. Nature Immunology, 2019. 20(1): p. 97-+.

34. Allie, S.R., et al., The establishment of resident memory B cells in the lung requires local antigen encounter. Nat Immunol, 2019. 20(1): p. 97–108.

35. Kwon, D.I., et al., Mucosal unadjuvanted booster vaccines elicit local IgA responses by conversion of pre-existing immunity in mice. Nat Immunol, 2025. 26(6): p. 908–919.

36. Liang, C., et al., Targeting heptad repeats and fusion peptide: nanoparticle vaccine elicits mucosal immune response against SARS-CoV-2 variants. J Nanobiotechnology, 2025. 23(1): p. 483.

37. Arroyo-Díaz, N.M., et al., Interferon-γ production by Tfh cells is required for CXCR3(+) pre-memory B cell differentiation and subsequent lung-resident memory B cell responses. Immunity, 2023. 56(10): p. 2358–2372.e5.

38. Risley, C.A., et al., Transcription factor T-bet regulates the maintenance and differentiation potential of lymph node and lung effector memory B cell subsets. Immunity, 2025. 58(7): p. 1706–1724 e6.

39. Adachi, Y., et al., Distinct germinal center selection at local sites shapes memory B cell response to viral escape. J Exp Med, 2015. 212(10): p. 1709–23.

40. Sampson, A.T., et al., Developing the next-generation of adenoviral vector vaccines. Hum Vaccin Immunother, 2025. 21(1): p. 2514356.

41. Elkashif, A., et al., Adenoviral vector-based platforms for developing effective vaccines to combat respiratory viral infections. Clin Transl Immunology, 2021. 10(10): p. e1345.

42. Chang, J., Adenovirus Vectors: Excellent Tools for Vaccine Development. Immune Netw, 2021. 21(1): p. e6.

43. Mendonca, S.A., et al., Adenoviral vector vaccine platforms in the SARS-CoV-2 pandemic. NPJ Vaccines, 2021. 6(1): p. 97.

44. Hassan, A.O., et al., A single intranasal dose of chimpanzee adenovirus-vectored vaccine protects against SARS-CoV-2 infection in rhesus macaques. Cell Rep Med, 2021. 2(4): p. 100230.

45. Hassan, A.O., et al., A Single-Dose Intranasal ChAd Vaccine Protects Upper and Lower Respiratory Tracts against SARS-CoV-2. Cell, 2020. 183(1): p. 169–184 e13.

46. Gabitzsch, E., et al., Dual-Antigen COVID-19 Vaccine Subcutaneous Prime Delivery With Oral Boosts Protects NHP Against SARS-CoV-2 Challenge. Front Immunol, 2021. 12: p. 729837.

47. Hatta, M., et al., Growth of H5N1 influenza A viruses in the upper respiratory tracts of mice. PLoS Pathog, 2007. 3(10): p. 1374–9.

48. Amalfitano, A., et al., Production and characterization of improved adenovirus vectors with the E1, E2b, and E3 genes deleted. J Virol, 1998. 72(2): p. 926–33.

49. Gabitzsch, E.S., et al., Control of SIV infection and subsequent induction of pandemic H1N1 immunity in rhesus macaques using an Ad5 [E1-, E2b-] vector platform. Vaccine, 2012. 30(50): p. 7265–70.

50. Morse, M.A., et al., Novel adenoviral vector induces T-cell responses despite anti-adenoviral neutralizing antibodies in colorectal cancer patients. Cancer Immunol Immunother, 2013. 62(8): p. 1293–301.

51. Jones, F.R., et al., Prevention of influenza virus shedding and protection from lethal H1N1 challenge using a consensus 2009 H1N1 HA and NA adenovirus vector vaccine. Vaccine, 2011. 29(40): p. 7020–6.

52. Risley, C.A., et al., T-bet expression marks a transcriptionally and functionally distinct population of memory B cells. Journal of Immunology, 2021. 206.

53. Zens, K.D., J.K. Chen, and D.L. Farber, Vaccine-generated lung tissue-resident memory T cells provide heterosubtypic protection to influenza infection. JCI Insight, 2016. 1(10).

54. McMaster, S.R., et al., Pulmonary antigen encounter regulates the establishment of tissue-resident CD8 memory T cells in the lung airways and parenchyma. Mucosal Immunol, 2018. 11(4): p. 1071–1078.

55. Wei, C.J., et al., Elicitation of broadly neutralizing influenza antibodies in animals with previous influenza exposure. Sci Transl Med, 2012. 4(147): p. 147ra114.

56. Ols, S., et al., Route of Vaccine Administration Alters Antigen Trafficking but Not Innate or Adaptive Immunity. Cell Rep, 2020. 30(12): p. 3964–3971 e7.

57. Kobayashi, Y., et al., Ultrasensitive protein-level detection for respiratory infectious viruses. Front Immunol, 2024. 15: p. 1445771.

58. Klinkhammer, J., et al., IFN-lambda prevents influenza virus spread from the upper airways to the lungs and limits virus transmission. Elife, 2018. 7.

59. Aeffner, F., B. Bolon, and I.C. Davis, Mouse Models of Acute Respiratory Distress Syndrome: A Review of Analytical Approaches, Pathologic Features, and Common Measurements. Toxicol Pathol, 2015. 43(8): p. 1074–92.

60. Chamanza, R. and J.A. Wright, A Review of the Comparative Anatomy, Histology, Physiology and Pathology of the Nasal Cavity of Rats, Mice, Dogs and Non-human Primates. Relevance to Inhalation Toxicology and Human Health Risk Assessment. J Comp Pathol, 2015. 153(4): p. 287–314.

61. Steppan, S., R. Adkins, and J. Anderson, Phylogeny and divergence-date estimates of rapid radiations in muroid rodents based on multiple nuclear genes. Syst Biol, 2004. 53(4): p. 533–53.

62. Neumann, K., et al., Molecular phylogeny of the Cricetinae subfamily based on the mitochondrial cytochrome b and 12S rRNA genes and the nuclear vWF gene. Mol Phylogenet Evol, 2006. 39(1): p. 135–48.

63. Imai, M., et al., Syrian hamsters as a small animal model for SARS-CoV-2 infection and countermeasure development. Proceedings of the National Academy of Sciences of the United States of America, 2020. 117(28): p. 16587–16595.

64. Braxton, A.M., et al., Hamsters as a Model of Severe Acute Respiratory Syndrome Coronavirus-2. Comparative Medicine, 2021. 71(5): p. 1–13.

65. Iwatsuki-Horimoto, K., et al., Syrian Hamster as an Animal Model for the Study of Human Influenza Virus Infection. Journal of Virology, 2018. 92(4).

66. Bouvier, N.M. and A.C. Lowen, Animal Models for Influenza Virus Pathogenesis and Transmission. Viruses, 2010. 2(8): p. 1530–1563.

67. Caceres, C.J., et al., Influenza antivirals and animal models. FEBS Open Bio, 2022. 12(6): p. 1142–1165.

68. Steppan, S.J. and J.J. Schenk, Muroid rodent phylogenetics: 900-species tree reveals increasing diversification rates. PLoS One, 2017. 12(8): p. e0183070.

69. Silva-Sanchez, A. and T.D. Randall, Role of iBALT in Respiratory Immunity. Curr Top Microbiol Immunol, 2020. 426: p. 21–43.

70. Gabitzsch, E.S., et al., Anti-tumor immunotherapy despite immunity to adenovirus using a novel adenoviral vector Ad5 [E1-, E2b-]-CEA. Cancer Immunol Immunother, 2010. 59(7): p. 1131–5.

71. Lee, C.M. and J.E. Oh, Resident Memory B Cells in Barrier Tissues. Front Immunol, 2022. 13: p. 953088.

72. Guillaume, S.M., et al., Lung B cells in ectopic germinal centers undergo affinity maturation. Proc Natl Acad Sci U S A, 2025. 122(14): p. e2416855122.

73. Schacht, S.S., et al., Activation and maturation of antigen-specific B cells in nonectopic lung infiltrates are independent of germinal center reactions in the draining lymph node. Cell Mol Immunol, 2025. 22(6): p. 612–627.

74. Son, Y.M., et al., Tissue-resident CD4(+) T helper cells assist the development of protective respiratory B and CD8(+) T cell memory responses. Sci Immunol, 2021. 6(55).

75. Swarnalekha, N., et al., T resident helper cells promote humoral responses in the lung. Sci Immunol, 2021. 6(55).

76. Krimpenfort, L.T., S.E. Degn, and B.A. Heesters, The follicular dendritic cell: At the germinal center of autoimmunity? Cell Rep, 2024. 43(3): p. 113869.

77. Shlomchik, M.J., W. Luo, and F. Weisel, Linking signaling and selection in the germinal center. Immunol Rev, 2019. 288(1): p. 49–63.

78. Allie, S.R. and T.D. Randall, Resident Memory B Cells. Viral Immunol, 2020. 33(4): p. 282–293.

79. Gregoire, C., et al., Viral infection engenders bona fide and bystander subsets of lung-resident memory B cells through a permissive mechanism. Immunity, 2022. 55(7): p. 1216–1233.e9.

80. Mathew, N.R., et al., Single-cell BCR and transcriptome analysis after influenza infection reveals spatiotemporal dynamics of antigen-specific B cells. Cell Rep, 2021. 35(12): p. 109286.

81. Ramirez, S.I., et al., Immunological memory diversity in the human upper airway. Nature, 2024. 632(8025): p. 630–636.

82. Tan, H.X., et al., Lung-resident memory B cells established after pulmonary influenza infection display distinct transcriptional and phenotypic profiles. Sci Immunol, 2022. 7(67): p. eabf5314.

83. Sicca, F., S. Neppelenbroek, and A. Huckriede, Effector mechanisms of influenza-specific antibodies: neutralization and beyond. Expert Rev Vaccines, 2018. 17(9): p. 785–795.

84. Chen, J., et al., Long term outcomes in survivors of epidemic Influenza A (H7N9) virus infection. Sci Rep, 2017. 7(1): p. 17275.

85. Keeler, S.P., et al., Influenza A Virus Infection Causes Chronic Lung Disease Linked to Sites of Active Viral RNA Remnants. J Immunol, 2018. 201(8): p. 2354–2368.

86. Wallick, C., et al., Impact of influenza infection on the short- and long-term health of patients with chronic obstructive pulmonary disease. J Med Econ, 2022. 25(1): p. 930–939.

87. Suri, C., et al., Interplay between Lung Diseases and Viral Infections: A Comprehensive Review. Microorganisms, 2024. 12(10).

88. Mantis, N.J., N. Rol, and B. Corthesy, Secretory IgA’s complex roles in immunity and mucosal homeostasis in the gut. Mucosal Immunol, 2011. 4(6): p. 603–11.

89. Brandtzaeg, P., Secretory IgA: Designed for Anti-Microbial Defense. Front Immunol, 2013. 4: p. 222.

90. Corthésy, B., Multi-faceted functions of secretory IgA at mucosal surfaces. Frontiers in Immunology, 2013. 4.

91. Binsker, U., et al., Immune exclusion by naturally acquired secretory IgA against pneumococcal pilus-1. J Clin Invest, 2020. 130(2): p. 927–941.

92. Zhang, G., et al., Nasal delivery of secretory IgA confers enhanced neutralizing activity against Omicron variants compared to its IgG counterpart. Mol Ther, 2025. 33(4): p. 1687–1700.

93. Quan, F.S., R.W. Compans, and S.M. Kang, Oral vaccination with inactivated influenza vaccine induces cross-protective immunity. Vaccine, 2012. 30(2): p. 180–8.

94. Li, Z., et al., Adjuvantation of Influenza Vaccines to Induce Cross-Protective Immunity. Vaccines (Basel), 2021. 9(2).

95. Bissett, C., et al., Systemic prime mucosal boost significantly increases protective efficacy of bivalent RSV influenza viral vectored vaccine. Npj Vaccines, 2024. 9(1).

96. Kim, J. and J. Chang, Cross-protective efficacy and safety of an adenovirus-based universal influenza vaccine expressing nucleoprotein, hemagglutinin, and the ectodomain of matrix protein 2. Vaccine, 2024. 42(15): p. 3505–3513.

97. Zhu, J., et al., The transcription factor T-bet is induced by multiple pathways and prevents an endogenous Th2 cell program during Th1 cell responses. Immunity, 2012. 37(4): p. 660–73.

98. Schultz, M.D., et al., A single intranasal administration of AdCOVID protects against SARS-CoV-2 infection in the upper and lower respiratory tracts. Hum Vaccin Immunother, 2022. 18(6): p. 2127292.

99. Jenkins, M.M., et al., Lung dendritic cells migrate to the spleen to prime long-lived TCF1(hi) memory CD8(+) T cell precursors after influenza infection. Sci Immunol, 2021. 6(63): p. eabg6895.

100. Hubner, R.H., et al., Standardized quantification of pulmonary fibrosis in histological samples. Biotechniques, 2008. 44(4): p. 507–11, 514-7.

101. Shu, B., et al., Multiplex Real-Time Reverse Transcription PCR for Influenza A Virus, Influenza B Virus, and Severe Acute Respiratory Syndrome Coronavirus 2. Emerg Infect Dis, 2021. 27(7): p. 1821–1830.

102. King, R.G., et al., Single-Dose Intranasal Administration of AdCOVID Elicits Systemic and Mucosal Immunity against SARS-CoV-2 and Fully Protects Mice from Lethal Challenge. Vaccines (Basel), 2021. 9(8).

103. Curtiss, M.L., et al., CXCR5 Signals Fine-Tune Dendritic Cell Transcription and Regulate T(H)2 Development. Vaccines (Basel), 2025. 13(9).

104. Beckett, D., E. Kovaleva, and P.J. Schatz, A minimal peptide substrate in biotin holoenzyme synthetase-catalyzed biotinylation. Protein Sci, 1999. 8(4): p. 921–9.

105. Bornholdt, Z.A. and B.V. Prasad, X-ray structure of NS1 from a highly pathogenic H5N1 influenza virus. Nature, 2008. 456(7224): p. 985–8.

