## Supplemental Figures and Tables for "Mucosal delivery of influenza antigens using a replication deficient adenovirus supports broadly reactive antibody responses and heterologous viral immunity in the respiratory tract of animals"

### Supplemental figures and legends

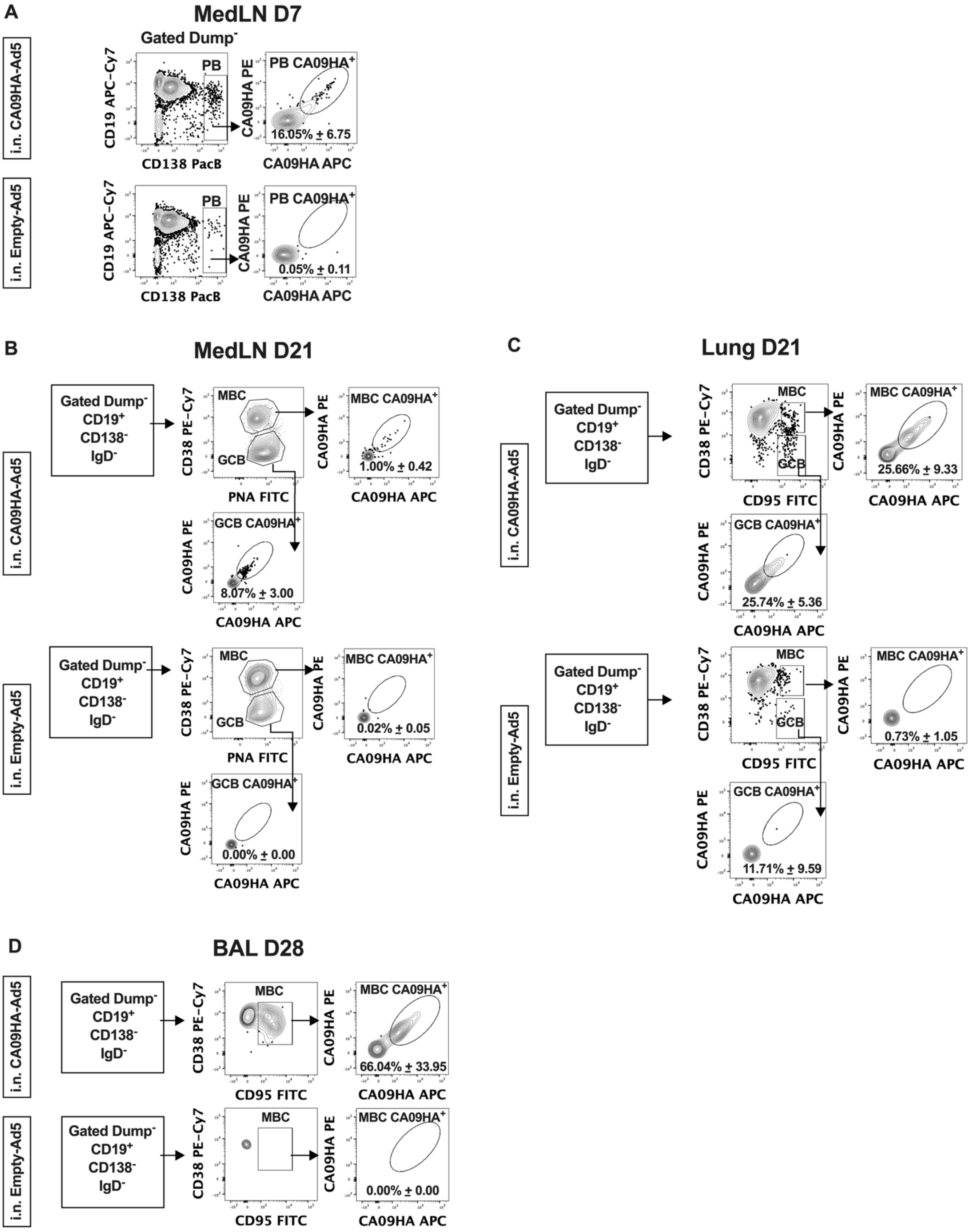

**Figure S1. Representative gating strategies for identification of CA09HA-specific B cell populations following CA09HA-Ad5 immunization. Related to Figure 1.**

Flow cytometry gating strategies to identify CA09HA$^+$ B cell subsets in the medLN and lung and BAL following i.n. immunization with CA09HA-Ad5 or Empty-Ad5.

(**A**) Identification of CD138$^+$CD19$^{+/lo}$ CA09HA$^+$ PB in MedLN on D7 post-immunization. Gated on live, Dump$^{neg}$ (CD64$^-$CD3$^-$) cells.

(**B**) Identification of CD38$^+$PNA$^{lo}$ CA09HA$^+$ MBC and CD38$^{lo}$PNA$^+$ GCB in MedLN on D21 post-immunization. Gated on live, Dump$^{neg}$ CD19$^+$CD138$^{neg}$IgD$^{neg}$ cells.

(**C**) Identification of CD38$^+$CD95$^+$ CA09HA$^+$ MBC and CD38$^{lo}$CD95$^+$ GCB in lung on D21 post-immunization. Gated on live, Dump$^{neg}$ CD19$^+$CD138$^{neg}$IgD$^{neg}$ cells.

(**D**) Identification of CD38$^+$CD95$^+$ CA09HA$^+$ MBC in BAL on D28 post-immunization. Gated on live, Dump$^{neg}$ CD19$^+$CD138$^{neg}$IgD$^{neg}$ cells.

Frequencies of each population reported as mean ± SD of 5 mice/group.

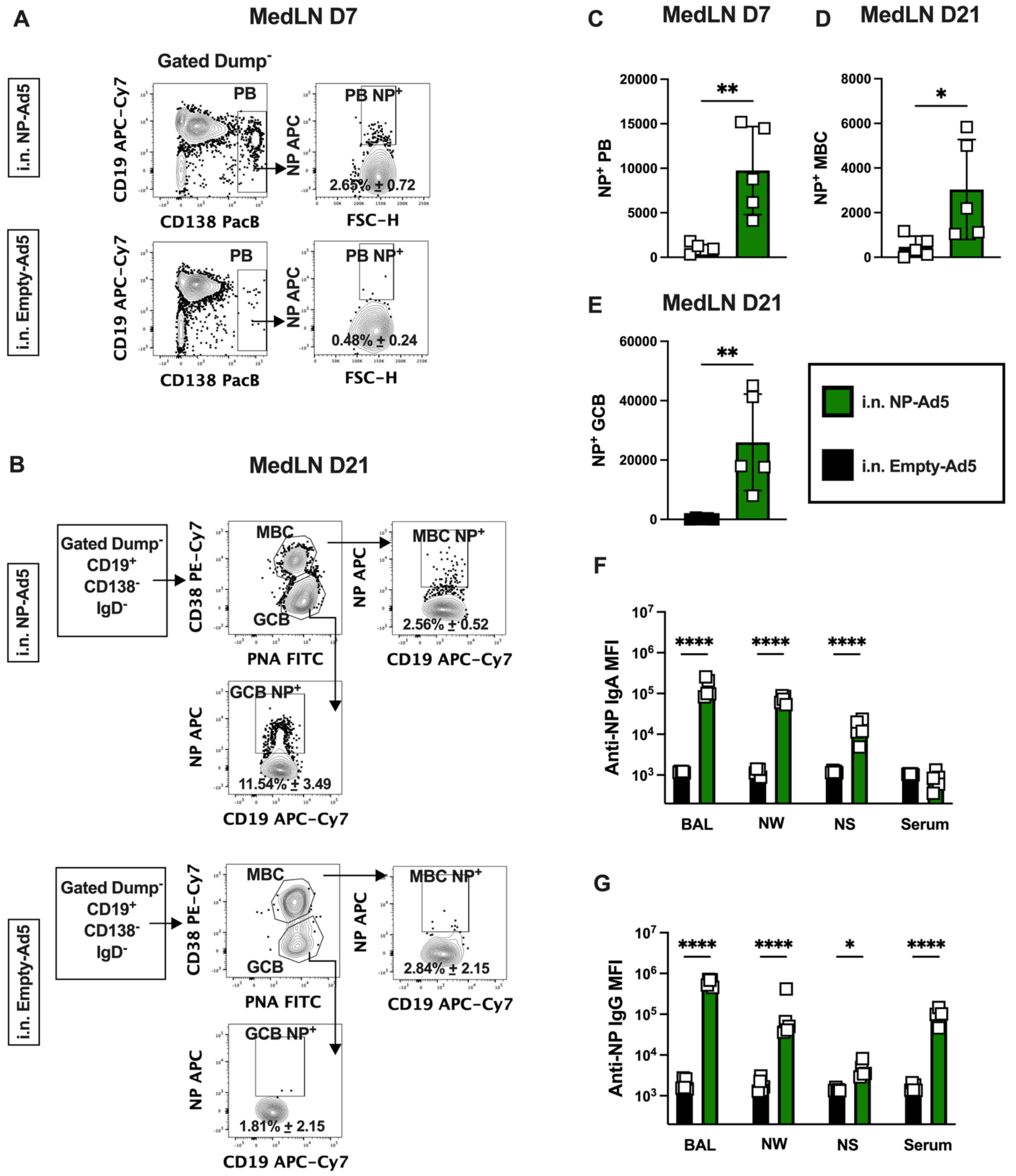


**Figure S2. Intranasal immunization with NP-Ad5 induces systemic and mucosal NP-specific B cell and Ab responses. Related to Figure 1.**

C57BL/6J (B6) mice (n=5 mice/group/timepoint/tissue sample) were immunized i.n. with Empty-Ad5 (black) or NP-Ad5 (green). Samples (MedLN, serum, BAL, NW, and NS) were collected at the indicated times and analyzed using flow cytometry (**A-E**) or CBA (**F-G**).

(**A-B**) Flow cytometry gating strategies to identify $NP^{+}$ PB (**A**), $NP^{+}$ MBC (**B**) and $NP^{+}$ GCB (**B**) in the medLN on D7 (**A**) and D21 (**B**). Frequencies of each population reported as mean ± SD.

(**C-E**) Number of D7 $NP^+$ PB (**C**), D21 $NP^+$ MBC (**D**) and D21 $NP^+$ GCB (**E**) from the MedLN. Data shown as mean ± SD of the medLN populations.
(**F-G**) NP-specific Ab responses in serum, BAL, NW, and NS samples collected at D28 post-immunization. NP-specific IgA (**F**) and IgG (**G**) levels were measured by CBA and reported as the mean ± SD of the log transformed MFI values of IgG or IgA Ab binding to NP-coupled microbeads.
Data analyzed by unpaired Mann–Whitney test (**C-E**) or two-way ANOVA with Tukey's multiple comparisons test (**F-G**). * $p < 0.05$, ** $p < 0.01$, *** $p < 0.001$, **** $p < 0.0001$.

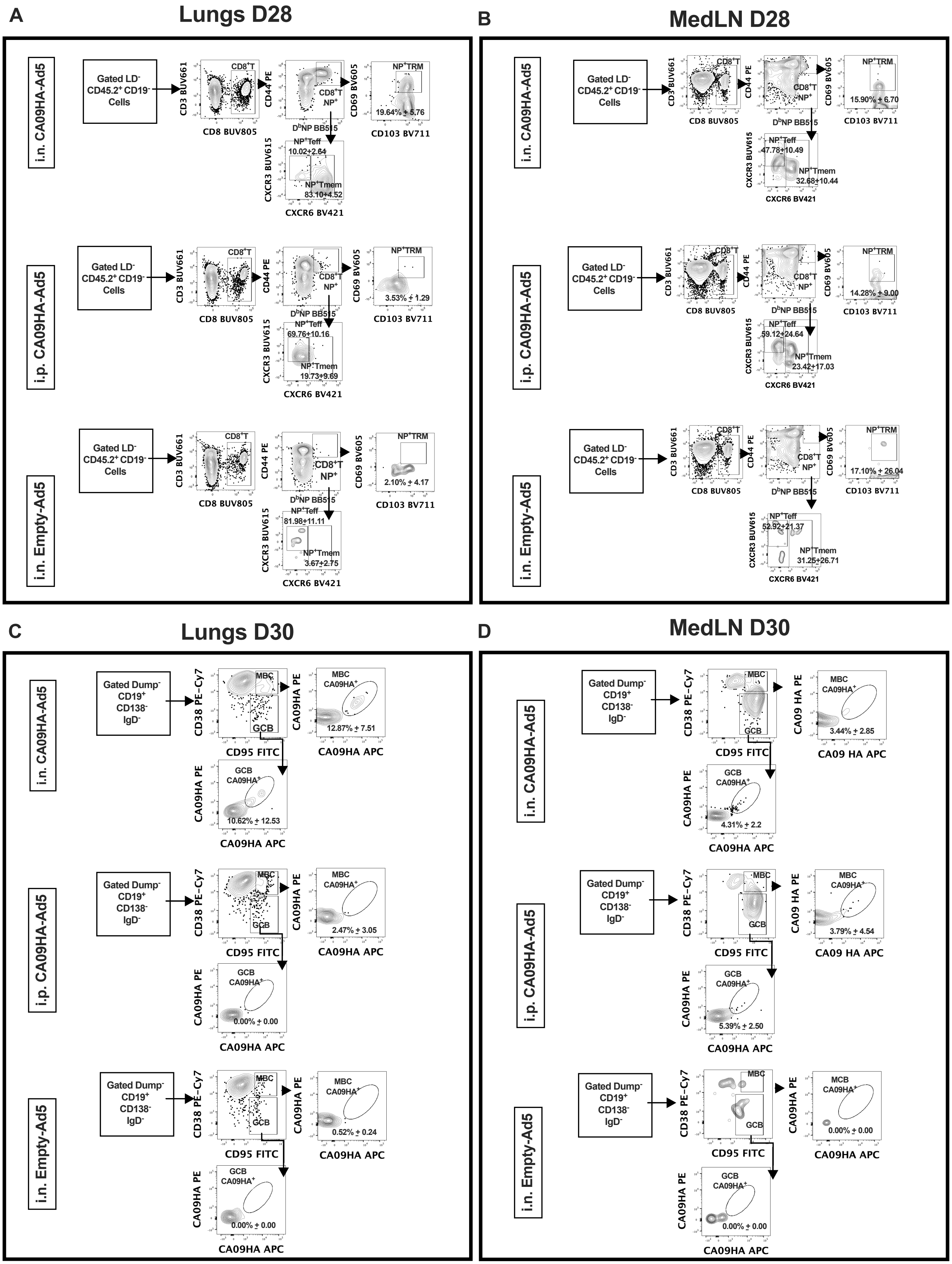

A
Lungs D28
B
MedLN D28
i.n. CA09HA-Ad5
i.p. CA09HA-Ad5
i.n. Empty-Ad5
Gated LD- CD45.2+ CD19- Cells
CD3 BUV661
CD8 BUV805
CD8+T
CD44 PE
DbNP BB515
CD8+T NP+
CD69 BV605
CD103 BV711
NP+TRM
CXCR3 BUV615
CXCR6 BV421
NP+Teff
NP+Tmem
19.64% ± 5.76
10.02±2.64
83.10±4.52
3.53% ± 1.29
69.76±10.16
19.73±9.69
2.10% ± 4.17
81.98±11.11
3.67±2.75
15.90% ± 6.70
47.78±10.49
32.68±10.44
14.28% ± 9.00
59.12±24.64
23.42±17.03
17.10% ± 26.04
52.92±21.37
31.25±26.71
C
Lungs D30
D
MedLN D30
Gated Dump- CD19+ CD138- IgD-
CD38 PE-Cy7
CD95 FITC
MBC
GCB
CA09HA PE
CA09HA APC
CA09 HA PE
CA09 HA APC
MBC CA09HA+
MCB CA09HA+
GCB CA09HA+
12.87% ± 7.51
10.62% ± 12.53
2.47% ± 3.05
0.00% ± 0.00
0.52% ± 0.24
3.44% ± 2.85
4.31% ± 2.2
3.79% ± 4.54
5.39% ± 2.50

**Figure S3. Intranasal HA vaccination uniquely elicits antigen-specific lung memory B and T cells. Related to Figure 4.**

Flow cytometry gating strategies to identify $NP^{+}$ CD8 T cell subsets (**A-B**) or $CA09HA^{+}$ B cell subsets (**C-D**) in the medLN and lung on D28 (**A-B**) or D30 (**C-D**) following i.n. immunization with Empty-Ad5 (all panels) or immunization with NP-Ad5 (**A-B**) or CA09HA-Ad5 (**C-D**) via the i.n. or i.p. route (n=4-5 mice/group).

(**A-B**) Identification of NP-specific $CD69^{+}CD103^{+}$ CD8 TRM, $CXCR3^{+}CXCR6^{neg}$ CD8 Teff and $CXCR6^{+}CXCR3^{+/-}$ CD8 Tmem on D28 post-immunization. Frequencies of the NP-specific CD8 T cell subsets in each tissue reported as mean ± SD.

(**C-D**) Identification of $CD38^{+}CD95^{+}$ $CA09HA^{+}$ MBC and $CD38^{lo}CD95^{+}$ GCB in lung (**C**) or MedLN (**D**) on D30 post-immunization. Gated on live, $Dump^{neg}$ $CD19^{+}CD138^{neg}IgD^{neg}$ cells. Frequencies of CA09HA-specific MBC and GCB in each tissue reported as mean ± SD.

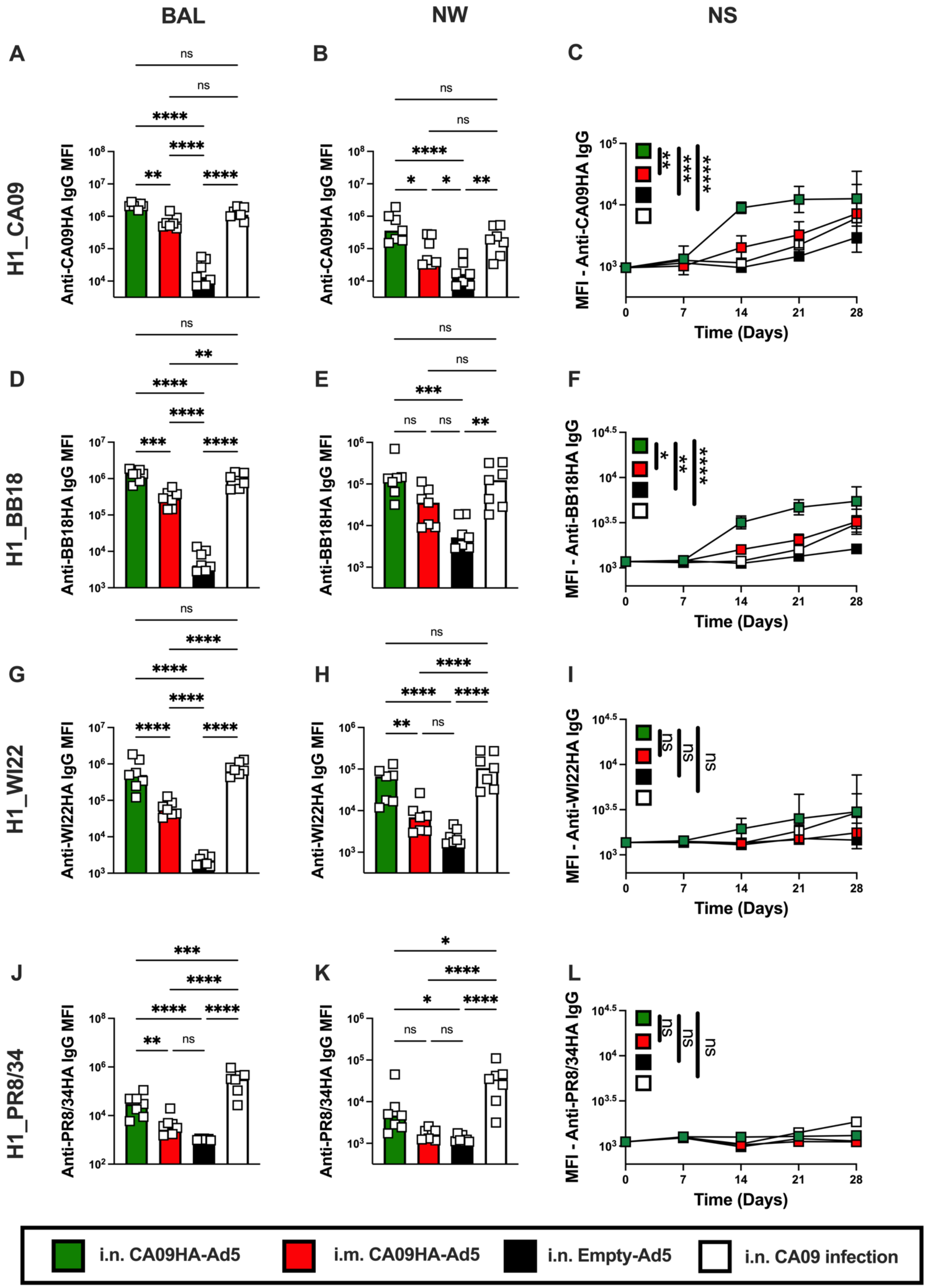
BAL
NW
NS
A
B
C
H1_CA09
Anti-CA09HA IgG MFI
MFI - Anti-CA09HA IgG
D
E
F
H1_BB18
Anti-BB18HA IgG MFI
MFI - Anti-BB18HA IgG
G
H
I
H1_WI22
Anti-WI22HA IgG MFI
MFI - Anti-WI22HA IgG
J
K
L
H1_PR8/34
Anti-PR8/34HA IgG MFI
MFI - Anti-PR8/34HA IgG
Time (Days)
i.n. CA09HA-Ad5
i.m. CA09HA-Ad5
i.n. Empty-Ad5
i.n. CA09 infection

1349

**Figure S4. Intranasal immunization with CA09HA-Ad5 supports development of IgG Abs with broad reactivity to H1_HA antigens. Related to Figure 5.**
B6 mice (n=7/group) were infected i.n. with 500 PFU H1_CA09 virus (white) or were immunized i.n. with Empty-Ad5 (black) or CA09HA-Ad5 via the i.n. (green) or i.m. (red) route. Samples were collected weekly (NS) or on D30 post-immunization/infection (BAL, NW). IgG Ab levels in samples were quantitated by CBA using microbeads coupled to recombinant H1_CA09HA, H1_BB18HA, H1_WI22HA, or H1_PR8/34HA antigens.
(**A-C**) Quantitation of H1_CA09HA-binding IgG in samples from BAL (**A**), NW (**B**), and NS (**C**).
(**D-F**) Quantitation of H1_BB18HA-binding IgG in samples from BAL (**D**), NW (**E**), and NS (**F**).
(**G-I**) Quantitation of H1_WI22HA-binding IgG in samples from BAL (**G**), NW (**H**), and NS (**I**).
(**J-L**) Quantitation of H1_PR8/34HA-binding IgG in samples from BAL (**J**), NW (**K**), and NS (**L**).
Data reported as mean ± SD (**A-B, D-E, G-H, J-K**) or mean ± SEM (**C, F, I, L**) of log transformed MFI values of Ab binding to the different HA-coupled microbeads.
Statistical analysis performed using two-way ANOVA with Tukey's multiple comparisons test (**A-B, D-E, G-H, J-K**) or using one-way ANOVA on AUC measurements (**C, F, I, L**). * $p < 0.05$, ** $p < 0.01$, *** $p < 0.001$, **** $p < 0.0001$.

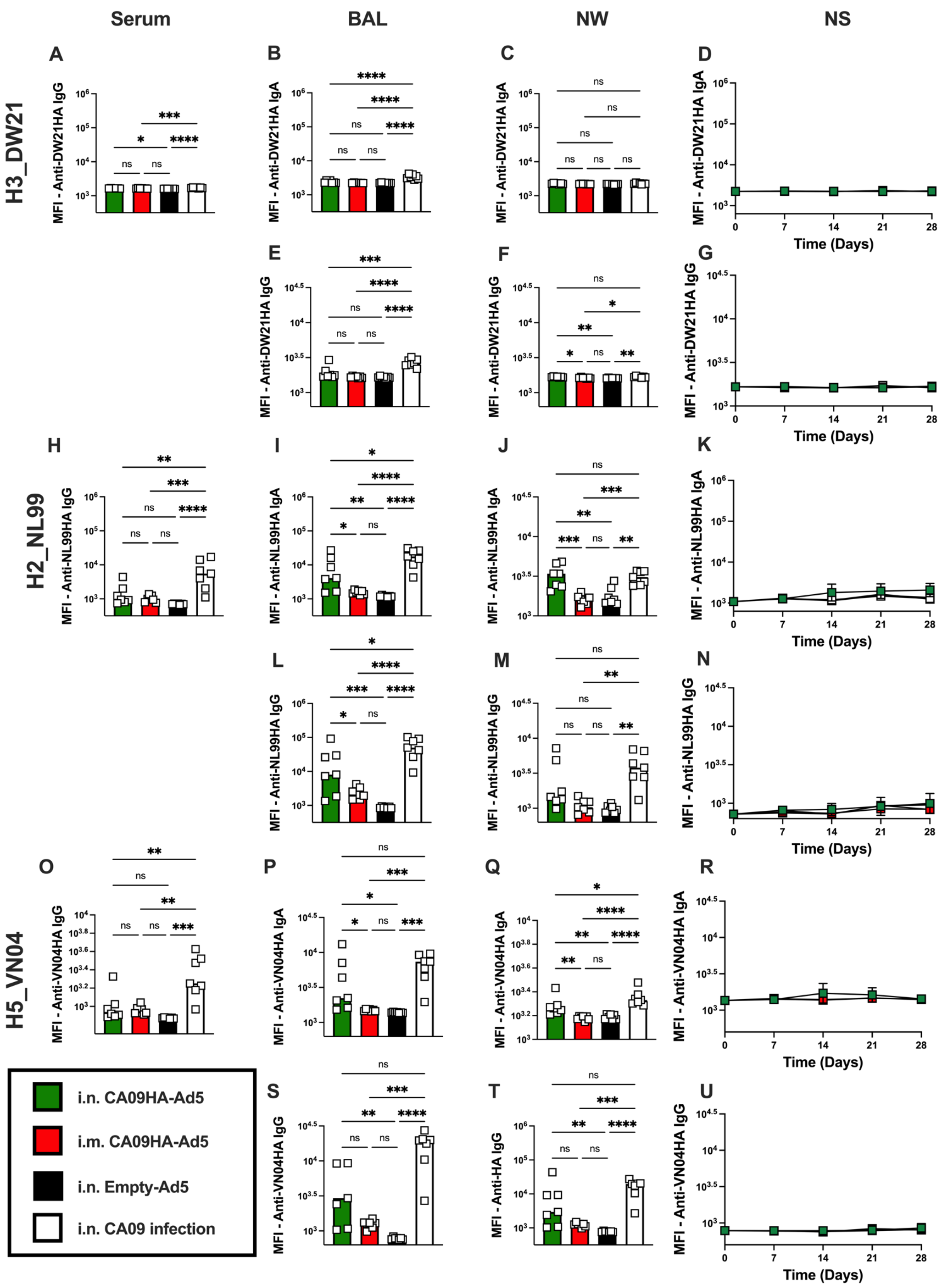

Serum
BAL
NW
NS
H3_DW21
H2_NL99
H5_VN04
A
B
C
D
E
F
G
H
I
J
K
L
M
N
O
P
Q
R
S
T
U
MFI - Anti-DW21HA IgG
MFI - Anti-DW21HA IgA
MFI - Anti-NL99HA IgG
MFI - Anti-NL99HA IgA
MFI - Anti-VN04HA IgG
MFI - Anti-VN04HA IgA
MFI - Anti-HA IgG
Time (Days)
i.n. CA09HA-Ad5
i.m. CA09HA-Ad5
i.n. Empty-Ad5
i.n. CA09 infection

**Figures S5. Intranasal immunization with CA09HA-Ad5 supports development of mucosal and systemic Abs that bind to avian influenza HA antigens. Related to Figure 5.**
B6 mice (n=7/group) were infected i.n. with 500 PFU H1_CA09 virus (white) or were immunized i.n. with Empty-Ad5 (black) or CA09HA-Ad5 via the i.n. (green) or i.m. (red) route. Samples were collected weekly (NS) or on D30 post-immunization/infection (serum, BAL, NW). IgG and IgA Ab levels in samples were quantitated by CBA using microbeads coupled to recombinant H3_DW21HA, H2_NL99HA, or H5_VN04HA antigens.
(**A-G**) Quantitation of serum IgG (**A**) binding to H3_DW21HA. Quantitation of IgA from BAL (**B**), NW (**C**), and NS (**D**) binding to H3_DW21HA. Quantitation of IgG from BAL (**E**), NW (**F**), and NS (**G**) binding to H3_DW21HA.
(**H-N**) Quantitation of serum IgG (**H**) binding to H2_NL99HA. Quantitation of IgA from BAL (**I**), NW (**J**), and NS (**K**) binding to H2_NL99HA. Quantitation of IgG from BAL (**L**), NW (**M**), and NS (**M**) binding to H2_NL99HA.
(**O-U**) Quantitation of serum IgG (**O**) binding to H5_VN04HA. Quantitation of IgA from BAL (**P**), NW (**Q**), and NS (**R**) binding to H5_VN04HA. Quantitation of IgG from BAL (**S**), NW (**T**), and NS (**U**) binding to H5_VN04HA.
Data reported as mean ± SD (**A-C, E-F, H-J, L-M, O-Q, S-T**) or mean ± SEM (**D, G, K, N, R, U**) of log transformed MFI values of Ab binding to the different HA-coupled microbeads.
Data evaluated using two-way ANOVA with Tukey's multiple comparisons test. *$p < 0.05$, ** $p < 0.01$, *** $p < 0.001$, **** $p < 0.0001$.

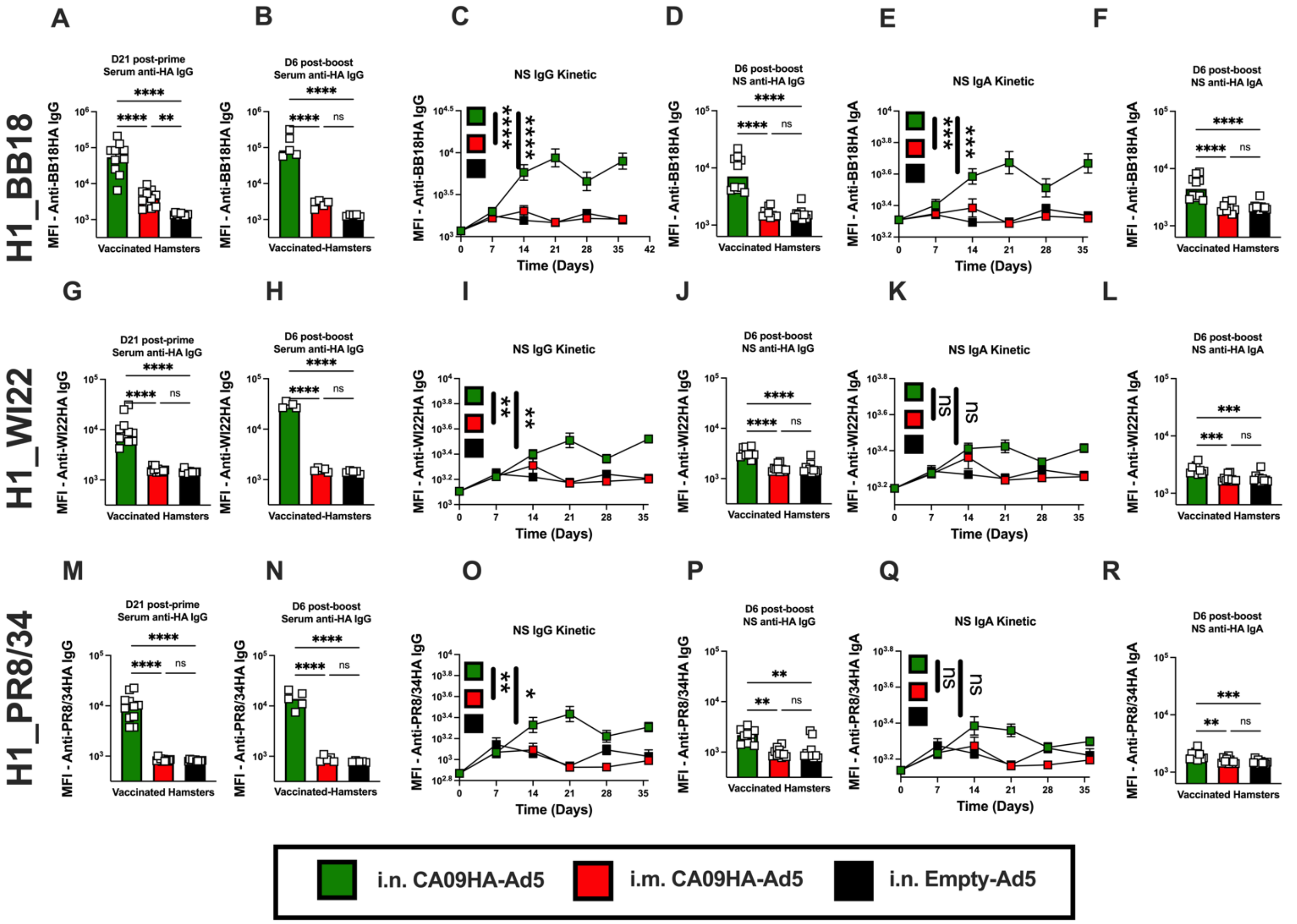


**Figure S6. Intranasal immunization of hamsters with CA09HA-Ad5 elicits broadly reactive Abs in serum and nasal secretions. Related to Figure 7.**
Hamsters (n=5-10/group/assay) were immunized i.n. with Empty-Ad5 (black) or immunized with CA09HA-Ad5 via the i.n. (green) or i.m. (red) route. Animals were boosted by the same route on D30 (i.n./i.n. or i.m./i.m.). NS samples were collected weekly from immunized animals (n=10/group). Serum was collected from immunized animals on D21 post-prime (n=10/group) and D6 post-boost (n=5/group). IgG and IgA Ab levels in samples were quantitated by CBA using microbeads coupled to recombinant H1_BB18HA, H1_WI22HA, or H1_PR8/34HA antigens.
(**A-F**) Quantitation of D21 post-prime (**A**) and D6 post-boost (**B**) serum IgG binding to H1_BB18HA. Quantitation of H1-BB18HA-binding IgG (**C-D**) and H1-BB18HA-binding IgA (**E-F**) from NS samples collected following prime and boost immunization.
(**G-L**) Quantitation of D21 post-prime (**G**) and D6 post-boost (**H**) serum IgG binding to H1_WI22HA. Quantitation of H1_WI22HA-binding IgG (**I-J**) and H1_WI22HA-binding IgA (**K-L**) from NS samples collected following prime and boost immunization.
(**M-R**) Quantitation of D21 post-prime (**M**) and D6 post-boost (**N**) serum IgG binding to H1_PR8/34HA. Quantitation of H1_PR8/34HA-binding IgG (**O-P**) and H1_PR8/34HA-binding IgA (**Q-R**) from NS samples collected following prime and boost immunization.
Data reported as mean ± SD or mean ± SEM (**C, E, I, K, O, Q**) of log transformed MFI values of Ab binding to the different HA-coupled microbeads. Statistical analyses performed using two-way ANOVA with Tukey's multiple comparisons test (**A-B, D, F, G-H, J, L, M-N, P, R**) or using one-way ANOVA on AUC measurements (**C, E, I, K, O, Q**). * $p < 0.05$, ** $p < 0.01$, *** $p < 0.001$, **** $p < 0.0001$.

### Table S1. Scoring rubric used to evaluate lungs from immunized and infected mice

| % Air-way affect-ed (0-4) | Airway severity (0-4) | Hyper plasia (0-3) | Air-way Score (0-11) | %Aveoli affected (0-4) | Alveolar Severity (0-4) | Type II pneumo-cyte hyper-plasia (0-3) | Asch-croft Score (0-8) | Alveol ar Score (0-19) | % vessels affected (0-4) | VascularPV Lesion Severity (0-3) | Necro-tizing Vascu-litis/ Thrombi (0-1) | Vas-cular Score (0-8) | Total Score (0-38) |
|---|---|---|---|---|---|---|---|---|---|---|---|---|---|
| 0= normal | 0= normal | 0= none | | 0= normal | 0= normal | 0=none | | | 0= normal | 0=none | 0= absent | | |
| 1= <10% | 1 = Mild peribronchitis/ bronchiolitis | 1 = <10% BEC hyperplasia | | 1= <10% | 1 = Mild peribronchiolar primarily mononuclear inflammatory infiltrates that occasionally extend into adj alveoar septa (1-3 cells thick) or alveolar spaces | 1 = Scattered type II pneumocyte hyperplasia affecting <10% of the section | | | 1= <10% | 1 = Multifocal perivascular edema and/or mild, primarily mononuclear perivascular inflammation (macrophages, lymphocytes +/- few scattered neutrophils | 1= present | | |
| 2= 10-25% | 2 = Mild to moderate mononuclear to mixed peribronchiolitis and/or lumens contain low numbers of inflammatory cells and/or multifocal single cell necrosis of airway epithelium | 2 = 10-25% BEC hyperplasia | | 2= 10-25% | 2 = Mild to moderate, mononuclear to mixed inflammation (>3 cells thick) expands alveolar septa or spaces and/or occasionally obscures normal septal architecture | 2 = Mild to moderate type II pneumocyte hyperplasia affecting 10-25% of the section (+/- atypical or multinucleated cells) | | | 2= 10-25% | 2 = Moderate mononuclear to mixed perivascular inflammation, edema or fibrin, with leukocytes occasionally transmigrating the vessel wall, and/or multifocal endotheliitis | | | |
| 3= 25-50% | 3 = Moderate to marked mixed peribronchitis or bronchiolitis, and/or large foci of bronchiolar epithelial necrosis, and or bronchiolar epithelium exhibit occasional atypical/ multinucleated syncytial cells | 3 = >25% BEC hyperplasia + syncytial cells | | 3= 25-50% | 3 = Moderate mixed interstitial inflammation, and/or alveolar damage characterized by type I pneumocyte necrosis/loss with replacement by hemorrhage, fibrin, edema, and/or necrotic debris (reminiscent of hyaline membranes), and or scattered atypical/syncytial cells | 3 = Widespread type II pneumocyte hyperplasia and/or atypical/ multinucleated syncytial cells affecting >25% of the section | | | 3= 25-50% | 3 = Severe mixed perivascular inflammation frequently expanding/replacing vessel wall and/or marked, frequent endotheliitis | | | |
| 4= >50% | 4 = Marked bronchiolitis and widespread epithelial necrosis +/- rupture of bronchiolar epithelium and/or bronchiolar epithelium exhibits frequent atypical/multinucleated syncytial cells | | | 4= >50% | 4 = Marked alveolar inflammation (mixed), alveolar septal damage (as described above) and loss of normal septal architecture with frequent atypical/syncytial cells | | | | 4= >50% | | | | |

#### Table S2. List of influenza antigens used in CBA and tetramer production

| Antigen | Virus strain | Accession # | Subtype classification | Bead code | Bead size | Bead peak |
|---|---|---|---|---|---|---|
| HA | A/mallard/Netherlands/12/1999 | AFM83145.1 | H2N9 | H2_NL1999 | 4 um | 2 |
| HA | A/Vietnam/1203/2004 | EPI1256205 | H5N1 | H5_VN2004 | 4 um | 3 |
| NP | A/Puerto Rico/8/1934 | NP_040982.1 | H1N1 | NP_PR8 | 4 um | 6 |
| NS1 | A/Puerto Rico/8/1934 | NP_040984.1 | H1N1 | NS1_PR8 | 4 um | 7 |
| HA | A/Puerto Rico/8/1934 | P03452.2 | H1N1 | H1_PR8 | 5 um | 8 |
| HA | A/California/07/2009 | ADB89281.1 | H1N1 | H1_CA2009 | 5 um | 2 |
| HA | A/Hawaii/70/2019 | QHN72768.1 | H1N1 | H1_HI2019 | 5 um | 3 |
| HA | A/Wisconsin/67/2022 | EPI15928538 | H1N1 | H1_WI2022 | 5 um | 4 |
| HA | A/Brisbane/02/2018 | OQ718997.1 | H1N1 | H1_BB2018 | 5 um | 5 |
| HA | A/Hong Kong/45/2019 | EPI588464 | H3N2 | H3_HK2019 | 5 um | 6 |
| HA | A/Darwin/9/2021 | EPI2233240 | H3N2 | H3_DW2021 | 5 um | 7 |
| HA | A/Singapore/INFIMH-16-0019/2016 | QQY97257.1 | H3N2 | H3_SG2016 | 5 um | 9 |

#### Table S3. Flow cytometry reagents used in this study

| Antibody/Tetramer | Company | Catalog # | Clone # | Fluorochrome | Dil (1:#) |
|---|---|---|---|---|---|
| CD19 | Biolegend | 115543 | 6D5 | BV785 | 200 |
| CD19 | Biolegend | 115558 | 6D5 | APC-Fire750 | 200 |
| CD138 | Biolegend | 142508 | 281-2 | BV421 | 200 |
| IgD | Biolegend | 405723 | 11-26c.2a | BV510 | 500 |
| CD38 | eBioscience | 25-0381-82 | 90 | PE-Cy7 | 400 |
| Fas(CD95) | BD | 554257 | Jo2 | FITC | 200 |
| PNA | Life Tech | L21409 | | AF488 | 400 |
| CD64 | Biolegend | 139308 | X54-5/7.1 | PerCP-Cy5.5 | 200 |
| CA09HA tetramer | In house | | | APC | 100 |
| CA09HA tetramer | In house | | | PE | 100 |
| NP | In house | | | APC | 100 |
| CD8α | BD | 612898 | 53-6.7 | BUV805 | 200 |
| CD8α | BD | 561967 | 53-6.7 | APC-Cy7 | 200 |
| CD4 | BD | 560782 | RM4-5 | V500 | 200 |
| CD3ε | Invitrogen | 45-0031-82 | 145-2C11 | PerCP-Cy5.5 | 200 |
| CD3ε | BD | 750638 | 145-2C11 | BUV661 | 200 |
| CD3ε | Biolegend | 155612 | KT3 | PacB | 100 |
| CD11b | BD Pharmingen | 550993 | M1/70 | PerCP-Cy5.5 | 200 |
| Live/Dead Aqua | ThermoFisher | L34965 | | V500 | 1000 |
| CD69 | Biolegend | 104529 | H1-2F3 | BV605 | 150 |
| CD103 | BD Horizon | 564320 | M290 | BV711 | 200 |
| CXCR3 | BD | 751457 | CXCR3-173 | BUV615 | 200 |
| CXCR6 | Biolegend | 151109 | SA051D1 | BV421 | 200 |
| CD45.2 | BD Horizon | 564616 | 104 | BUV395 | 200 |
| CD62L | BD | 560514 | MEL-14 | APC-Cy7 | 400 |
| CD8 NP366-374 H2D(b) tetramer | NIH | 78600 | | BB515 | 100 |
| CD8 NP366-374 H2D(b) tetramer | NIH | 77852 | | APC | 100 |
| 7AAD | Biolegend | 420403 | | PerCP-Cy5.5 | 1000 |
